# The rapid diversification of Boletales is linked to Early Eocene and Mid-Miocene Climatic Optima

**DOI:** 10.1101/2023.10.24.563795

**Authors:** Gang Wu, Kui Wu, Roy E. Halling, Egon Horak, Jianping Xu, Guang-Mei Li, Serena Lee, Lorenzo Pecoraro, Roberto Flores Arzu, Sydney T. Ndolo Ebika, Souhila Aouali, Anna Maria Persiani, Nourou S. Yorou, Xin Xu, Bang Feng, Yan-Chun Li, Zhu-Liang Yang

**Affiliations:** CAS Key Laboratory for Plant Diversity and Biogeography of East Asia, Kunming Institute of Botany, Chinese Academy of Sciences, Kunming 650201, Yunnan, China; Yunnan Key Laboratory for Fungal Diversity and Green Development, Kunming 650201, Yunnan, China; Institute of Systematic Botany, New York Botanical Garden, 2900 Southern Blvd., Bronx, NY 10458-5126, USA; Schlossfeld 17, Innsbruck A-6020, Austria; Department of Biology, McMaster University, Hamilton, ON L8S 4K1, Canada; Singapore Botanic Gardens, 1 Cluny Road, Singapore 259569; School of Pharmaceutical Science and Technology, Tianjin University, Tianjin 300072, China; Departamento de Microbiología, Escuela de Química Biológica, Facultad de Ciencias Químicas y Farmacia, Universidad de San Carlos de Guatemala, Zona 12, Ciudad de Guatemala, Guatemala; Faculty of Sciences and Techniques, Marien Ngouabi University, BP 69 Brazzaville, Republic of Congo; Initiative des Champignons et des Plantes du Congo, BP 2300, Brazzaville, Republic of Congo; Forest Pathology and Mycology Laboratory, Forest Research National Institute, B.P. 37, Cheraga, 16000, Algiers, Algeria; Fungal Biodiversity Laboratory, Department of Environmental Biology, Sapienza University of Rome, P.le A. Moro 5, 00185, Rome, Italy; Research Unit Tropical Mycology and Plants-Soil Fungi Interactions, Faculty of Agronomy, University of Parakou, BP 123 Parakou, Benin

**Keywords:** Agaricomycetes, ectomycorrhizal fungi, paleoclimate, phylogenomic analysis, spatiotemporal evolution, species diversity, symbiosis

## Abstract

- Investigating the mechanisms that underpin the diversity and distribution patterns of species is fundamental in ecology and evolution. However, the study of fungi, particularly the ectomycorrhizal group, has been relatively constrained in this field.
- We conducted a high-resolution phylogenomic analysis of Boletales, an ecologically and economically significant group of fungi, covering 83 genera across 15 families. We subsequently investigated its evolutionary history using sequences at four genes obtained from 984 species across 121 genera within 15 families.
- The findings unveiled that Boletales likely originated in Early Jurassic and underwent two remarkable episodes of rapid diversification, commencing in early Eocene (∼54 Mya) and early Miocene (∼17 Mya) epochs. The notable surges were predominantly driven by ectomycorrhizal clades, with a specific emphasis on East Asia and North America. These expansions were strongly correlated with the warm-humid paleoclimates during the Early Eocene Climatic Optimum and Mid-Miocene Climatic Optimum, as well as the rapid expansion of Fagales and Pinaceae hosts.
- This study provides novel insights into the spatiotemporal evolution of fungi, highlighting the synergistic impact of abiotic factors, such as warm and humid paleoclimates, and the biotic factor of rapid diversification of host plants on the fungal diversification.

## INTRODUCTION

Understanding the drivers of species diversity and its distribution pattern is among the fundamental goals in ecology and evolution (Chazot *et al*., 2021; Jin *et al*., 2021). Many researchers have made significant progress in understanding such drivers on alpine flora (Ding *et al*., 2020), bamboos (Guo *et al*., 2020), birches (Yang *et al*., 2022), oaks (Zhou *et al*., 2022), pines (Jin *et al*., 2021) and rosids (Ellepola *et al*., 2022) in plants; and butterflies (Chazot *et al*., 2021), birds (Aves) (Prum *et al*., 2015) and frogs (Ellepola *et al*., 2022) in animals. Their investigations revealed significant influences of Cenozoic climatic and geological events, such as the fluctuations of global temperature and aridity, mountains uplifting, on species diversification.

However, partially due to the limited fossil record and a lack of comprehensive phylogenies for many groups of fungi, there have been only a few such studies in class Agaricomycetes. In one representative study, a rapid radiation of this class was found in the Jurassic period, followed by potential mass extinctions, specific adaptive radiations, and morphological diversification of fruiting bodies during the Cretaceous and Paleogene periods (Varga *et al*., 2019). Another study by Sánchez-García *et al*. (2020) found a significant correlation between the pileate-stipitate form and the rapid diversification rates of Agaricomycetes, independent of nutritional mode. However, a recent finding by Sato (2023) contradicts this by suggesting that the evolution of ectomycorrhizal symbiosis in the Late Cretaceous was a key factor in the explosive diversification of Agaricomycetes.

Boletales is one of the most species-rich orders in the Agaricomycetes, with around 3000 described species according to Mycobank and Fungal Names databases. The suborders Boletineae and Suillineae play a significant role in the diversity of this order, making up approximately 2000 and 400 described species respectively. In terms of distribution, East Asia has been suggested as the center of existing species diversity for certain bolete groups like *Boletus* and *Strobilomyces* (Feng et al., 2012; Han et al., 2018). This order, especially its family Boletaceae, is also well known for their delicious edible mushrooms such as the worldwide-famous porcini (*Boletus edulis* complex) and the regionally popular *Lanmaoa asiatica*, *Butyriboletus roseoflavus*, *Suillus luteus*, etc. In addition to their culinary value, Boletales species perform crucial functions as brown-rot and ectomycorrhizal fungi that regulate the carbon and nitrogen cycles of forest ecosystems (Lindahl & Tunlid, 2015; Wu *et al*., 2022). A recent study based on 100 species of Boletales using 87 single-copy genes revealed that Boletales diversified rapidly, with signatures of co-evolution with angiosperms (particularly Fagaceae), in young ectomycorrhizal lineages (Sato & Toju, 2019).

However, the evolutionary impacts of the paleo-climatic and geological changes affecting Boletales in the Cenozoic remain unclear, despite the remarkable magnitude and extent of those changes during that period (Zachos *et al*., 2001). The increasing availability of Boletales specimens from many parts of the world with associated DNA sequence information, including whole-genome sequences, make Boletales an ideal group to investigate how evolutionary and ecological factors have impacted the spatiotemporal evolution in fungi.

Through the utilization of genomic and four-gene sequence data, this study aims to elucidate the intricate diversification history of the Boletales order and investigate its potential correlation with paleoclimatic changes and the diversification of host plants, exploring whether there is a discernible relationship. Additionally, we aim to determine the relative significance of these environmental and biotic factors in shaping the diversification of Boletales. Furthermore, our research delves into the genetic reticulate evolution and morphological adaptation that may influence the diversification of the Boletales order, providing valuable insights into its evolutionary journey.

## MATERIALS & METHODS

### Taxon sampling and necessary morphological study

To efficiently retrieve and economically obtain genome data, we utilized deep genome skimming (Dentinger *et al*., 2016; Guo *et al*., 2020) and newly sequenced a total of 64 species from 63 genera belonging to five families within the Boletales order, as well as one outgroup species from the Lepidostromatales order. To perform four-gene phylogenetic analyses, we collected a total of 984 samples from 984 phylogenetic species, belonging to 119 genera in 16 families within the Boletales. Some of the samples used in this study were collected from different regions in China and deposited at the Cryptogamic Herbarium (HKAS) of the Kunming Institute of Botany, Chinese Academy of Sciences. Others were loaned from various herbaria, including GDGM from Guangdong Province of China, TNS from Japan, NY from the USA, ZT from Switzerland, ROHB from Italy, SING from Singapore, MICG from Guatemala, HICPC from Republic of Congo. Data of other samples were retrieved from GenBank Nucleotide database. The information about all analyzed specimens is presented in Supporting Information Table S1 and S2. In order to better confirm and delimit species, we investigated necessary macro- and micro-morphological features following protocols described in Wu *et al*. (2016b), and Wu *et al*. (2014) for SEMs of basidiospores. The processes of DNA extraction, sanger sequencing and next-generation-sequencing were described in Supporting Information-Methods S1.

### Data assembly of genome skimming and determination of single-copy orthologues

To ensure the quality of the next-generation-sequencing reads, we first performed quality control using *fastp* (Chen *et al*., 2018). The reads were then assembled using *SPAdes* (Bankevich *et al*., 2012) with default settings. Subsequently, we conducted BUSCO analysis for each new assembly, as well as for the published genomes included in this study, using the boletales_odb10 database (Simão *et al*., 2015). The BUSCO results are shown in Supporting Information Table S1. To identify single-copy orthologues from the genome skimming assemblies as many as possible, we screened the shared protein sequences from the output files named *single_copy_busco_sequences* generated by the BUSCO analyses for each sample.

### Species determination for phylogenetic analyses

Apart from the species of Boletales with new genomic assemblies, representative orders of Basidiomycota and allied outgroups with published genomes were included in the phylogenomic analysis. The published genomes were chosen from JGI MycoCosm (https://mycocosm.jgi.doe.gov/mycocosm/home).

To expand the number of included Boletales species for four-gene phylogenetic analysis, we downloaded all available sequences of Boletales species from the GenBank Nucleotide database using the keyword “Boletales” on 3 March 2023. We then used a custom Python script to screen sequences of four DNA fragments (nrLSU, *TEF1*, *RPB1*, *RPB2*) and mapped them to shared voucher specimens using the keywords “strain”, “specimen_voucher”, “isolate” and “clone” in GenBank (.gb) files. We determined their distribution information using the keyword “country” in GenBank files. We subsequently built a phylogenetic tree using specimens with at least two of the above four DNA fragments.

Species delimitation in Boletales was based on the following criteria: 1) undelimited phylogenetic species were determined with a < 97% DNA sequence identity threshold between sibling species, 2) well-studied taxa were based on previous taxonomic and phylogenetic publications (Binder & Hibbett, 2006; Binder *et al*., 2010; Halling *et al*., 2012; Wilson *et al*., 2012; Nuhn *et al*., 2013; Halling *et al*., 2015; Henkel *et al*., 2016; Wu *et al*., 2016a; Wu *et al*., 2016b; Zeng *et al*., 2016; Zeng *et al*., 2017; Zeng *et al*., 2018; Chai *et al*., 2019; Kuo & Ortiz-Santana, 2020; Li & Yang, 2021; Magnago *et al*., 2022). During the determination, a small proportion of those known species had genetic variation of less than 3% between sister species. For species with newly generated sequences in this study, we identified them based on molecular and morphological evidence. The processes of sequence alignment were described in Supporting Information-Methods S1

### Phylogenomic and multi-gene phylogenetic analyses

The best-fitting model for each nucleotide and protein sequences was evaluated in ModelTest-NG with default settings (Darriba *et al*., 2019). For the phylogenetic analysis, two steps were conducted, including an initial phylogenomic analysis and a subsequent four-gene phylogenetic analysis. To minimize incongruences among the screened gene sequences (typically translated protein sequences) in phylogenomic analysis, we examined the incongruence between likelihood-based signal (measured by the difference in gene-wise log-likelihood score or ΔGLS) and quartet-based topological signal (measured by the difference in gene-wise quartet score or ΔGQS) for every protein, by following the method of Shen *et al*. (2021). *IQ-TREE* v2.2.0.3 (Nguyen *et al*., 2014) was used to produce gene trees, and the species tree was constructed using *ASTRAL* 5.7.8 (Mirarab *et al*., 2014).

To balance species coverage and the computational resources required, we generated two genomic datasets - D1 and D2. The D1 dataset included congruent gene sequences shared by 80% (considering a broader range of species and a greater number of genes) of the included species, which was used to reconstruct a high-resolution phylogenetic topology. The D2 dataset, on the other hand, included congruent gene sequences shared by 90% (considering a broader range of species and shorter computational time) of the included species, which was used to infer divergence times. Based on the D1 dataset, a phylogenetic tree was reconstructed using maximum likelihood in *IQ-TREE* v2.2.0.3 (Nguyen *et al*., 2014) with default settings, except -b was set to 100. Based on the D2 dataset, a maximum likelihood analysis was conducted by *RAxML* v8.2.11 (Stamatakis, 2014) with default settings, except the substitutional model set as PROTGAMMALG, and the number of bootstrapping was set to 100. The phylogenomic framework of Boletales based on D1 dataset was used as a constraint topology for the subsequent four-gene phylogenetic analysis. When analyzing, the constraint topology was pre-specified, and four related outgroup taxa were selected: *Sulzbacheromyces yunnanensis*, *Piloderma olivaceum*, *Plicaturopsis* sp., and *Agaricus bisporus* var. *bisporus*. Maximum likelihood analysis was carried out in *RAxML* v8.2.11 (Stamatakis, 2014), with default settings, except the substitutional model set as GTRGAMMAI, and statistical support was obtained using nonparametric bootstrapping with 1000 replicates.

### Divergence time estimation

The analysis of divergence time based on the D2 dataset were conducted using MCMCTree method implemented in *PAML* v.4.9j (Yang, 2007) by following their tutorial (http://abacus.gene.ucl.ac.uk/software/MCMCtree.Tutorials.pdf). The analysis utilized the independent-rates clock model, a WAG substitution model, and approximate likelihood calculation. The specific parameter settings used were as follows: *burnin* = 2000000, *sampfreq* = 100, and *nsample* = 100000. Five fossils were selected referring to previous studies (Varga *et al*., 2019), namely *Quatsinoporites cranhamii* (crown age of Hymenochaetales: 127-250 Mya, uniform distribution) (Smith *et al*., 2004), *Palaeoagaricites antiquus* (crown age of Agaricales: 105-210 Mya, uniform distribution) (Poinar & Buckley, 2007), *Rhizopogon*/*Suillus* ECM (crown age of Suillineae: 50-100 Mya, uniform distribution) (LePage *et al*., 1997), *Trametites eocenicus* (crown age of *Trametes*: 45-90 Mya, uniform distribution) (Knobloch & Kotlaba, 1994), and *Ganodermites lybicus* (crown age of *Ganoderma*: 18-50 Mya, uniform distribution) (Fleischmann *et al*., 2007). We subsequently employed the ages (a 95% confidence interval) of the backbone nodes of Boletales estimated by MCMCTree as input parameters for the divergence time analysis based on four-gene sequences. This analysis employed the *RelTime*-ML method implemented in *MEGA* 11 (Kumar *et al*., 2018) with default settings.

### Diversification analyses

We estimated diversification rates using the calibrated four-gene phylogenetic tree excluding outgroups. A lineage through time (LTT) plot was generated using *ape* v5.7.1 (Paradis & Schliep, 2018). To investigate the process of species diversification in different regions, we analyzed the accumulation of lineages through time using the data generated by *ltt.plot.coords* implemented in *ape* v5.7.1. For this analysis, the samples were separated into nine geographic regions used in previous biogeographical studies of boletes (Feng *et al*., 2012; Han *et al*., 2018): East Asia, Southeast Asia, South Asia, Australasia, North America, Central America, South America, Europe, and Africa. The location information of the included species was collected from various sources, including GenBank Nucleotide database and loaned or newly collected specimens (Supporting Information Table S2).

We firstly used Bayesian Analysis of Macroevolutionary Mixtures (BAMM v2.5.0) (Rabosky, 2014) to analyze shifts in diversification by following the work of Sato et al. (2017) and Sato and Toju (2019). In order to determine the sampling fraction, we initially adopted the approach proposed by Sato and Toju (2019) to estimate the number of Boletales species, considering the current lack of clarity regarding the total number of species within this order. We downloaded the nucleotide sequences of rRNA internal transcribed spacer (ITS) regions of Boletales from the NCBI website (http://www.ncbi.nih.gov). Sequences with more than four ambiguous bases (N) or shorter than 100 bp, or obtained from environmental (metabarcoding) samples were excluded. The remaining sequences were clustered into operational taxonomic units (OTUs) at a threshold of sequence identity 97% using *vsearch* software. Following Sato and Toju (2019), we used the “*specpool*” and “*specaccum*” functions in the R package *vegan* 2.6.2 to estimate the global richness of Boletales OTUs, which was estimated to 3227. The sampling fraction of Boletales was calculated by dividing the number of Boletales species (984) included in the phylogenetic analysis by the estimated global OTU richness (3227); the result was 0.305.

To ensure that the effective sizes of *postburn$N_shifts* and *postburn$logLik* were both over 200, we performed BAMM with 20,000,000 Markov chain Monte Carlo (MCMC) generations. The sampling fraction was set to 0.305 and the priors were calculated by the “*setBAMMpriors*” function implemented in the R package *BAMMtools* v2.1.10 (Rabosky *et al*., 2014). We used the “*getBestShiftConfiguration*” function to extract the rate shift configuration with the highest posterior probability.

The “*plot.bammdata*” function was used to plot the results. We used the “*plotRateThroughTime*” function to plot speciation, extinction, and net diversification rates through time for the following clades: Boletales, Boletineae (Boletaceae), Paxillineae, Boletinellineae (Boletinellaceae), Sclerodermatineae, Suillineae, Coniophorineae, and Tapinellineae, circumscribed by Binder and Hibbett (2006), as well as *Austroboletoideae*, *Boletoideae*, *Chalciporoideae*, *Leccinoideae*, *Pseudoboletoideae* subfam. nov, *Suillelloideae* subfam. nov. *Xerocomoideae*, and *Zangioideae* in family Boletaceae, circumscribed by Wu *et al*. (2014) and this study together. We also plotted the ectomycorrhizal group using the Boletineae, Paxillineae, Boletinellineae, Sclerodermatineae, and Suillineae and the saprotrophic group with Coniophorineae and Tapinellineae. To compare the differences in diversification rates among different clades, we used the R package *vioplot* v0.3.7 (https://rdrr.io/cran/vioplot/) to plot them together.

In light of recent criticisms regarding BAMM’s modeling of rate-shifts on extinct lineages (Moore *et al*., 2016), we have opted to incorporate additional Bayesian-based estimations of episodic diversification rate and branch-specific diversification rate using the Birth-Death models in RevBayes v3.3 (Höhna *et al*., 2016; Condamine *et al*., 2018). This approach ensures consistent modeling of rate shifts on extinct lineages.

We followed the tutorials available at https://revbayes.github.io/tutorials/ to guide our implementation. Meanwhile, we estimated maximum likelihood speciation and extinction rates, along with the shift times, using episodic Birth-Death model implemented in the R-package *TreePar* v3.3 (Stadler, 2011). We employed its “*bd.shifts.optim*” function to estimate discrete changes in speciation and extinction rates, as well as mass extinction events, in incompletely sampled phylogeny. We searched the entire timespan of the phylogeny at 1-million-year intervals for the likelihood of shifts in diversification and up to eight mass extinction events. To account for under-sampling in the tree, we set the sampling parameter to 0.305 (984/3227). The processes of dispersal analyses and phylogenetic network analyses were described in Supporting Information-Methods S1

### Correlation analyses between potential factors and the diversification of Boletales

We conducted correlation analyses to assess the potential factors, namely paleo-temperature, paleo-aridity, Fagales host, and Pinaceae host, that might correlate the diversification process of Boletales. To achieve this, we performed the estimates of environmental-dependent Speciation & Extinction Rates in *RevBayes* v3.3 (Höhna *et al*., 2016; Condamine *et al*., 2018) by following the tutorials available at https://revbayes.github.io/tutorials/.

We firstly collected original data of global paleo-temperature since 65 million years ago (Mya) from Zachos *et al*. (2008) and the simulated data of global paleo-aridity index since 65 Mya from Hagen *et al*. (2021).

To estimate the diversification rates of Fagales host plants, we utilized the phylogenetic tree reconstructed by Xing *et al*. (2014). We applied a similar procedure to Boletales using *RevBayes* v3.3 (Höhna *et al*., 2016) to calculate the diversification rates for Fagales. The sampling parameter was set to 0.423 (579/1370), with the total number of Fagales species obtained from Xing *et al*. (2014).

As for Pinaceae host plants, due to the lack of an appropriate phylogenetic tree, we assembled available transcriptomic data of the plants such as *Pinus* spp., *Picea* spp., *Tsuga* spp., *Cathaya argyrophylla*, *Pseudotsuga menziesii*, *Larix gmelinii*, *Nothotsuga longibracteata*, *Pseudolarix amabilis*, *Keteleeria evelyniana*, *Cedrus deodara*, and outgroups *Platycladus orientalis* and *Araucaria cunninghamii*. The data sources included the studies by Ran *et al*. (2018), Shen *et al*. (2019), Feng *et al*. (2021), and Jin *et al*. (2021). We reconstructed a high-resolution phylogenetic tree for Pinaceae using these transcriptomic data. To identify shared common coding DNA sequences (CDS) from the datasets, we utilized 1662 CDS from the *Pinus* data of Jin et al. (2021) as the initial reference sequences to map the CDS shared with the *Tsuga* CDS in the Feng et al. (2021) dataset. Then, we used the shared CDS between *Pinus* and *Tsuga* to map the shared CDS with the *Picea* data of Shen et al. (2019), as well as the remaining Pinaceae and outgroups CDS from Ran et al. (2018) using *Geneious* v2020.0.5. The diversification analysis for Pinaceae followed the same methodology as that for Boletales by using *RevBayes* v3.3 (Höhna *et al*., 2016). The sampling parameter was set to 0.635 (146/230), with the total number of Pinaceae species obtained from Du *et al*. (2020).

In the correlation analyses, we divided the paleo-temperature, simulated paleo-aridity index, and net diversification rates of Fagales and Pinaceae into 100 bins spanning from 65 Mya to present. By following the tutorials, we respectively applied four environmentally-dependent diversification models, namely GMRF (Gaussian Markov Random Field), HSMRF (Horseshoe Markov Random Field), UC (uncorrelated) and *fixed*, and two MCMC runs were conducted with 100000 iterations for each in this analysis (Höhna *et al*., 2016; Palazzesi *et al*., 2022)

### Ancestral character reconstruction of key morphologies

To investigate the potential factors driving the evolution of key morphological characters of Boletales, we focused on three taxonomically important morphological traits: the presence of the universal veil on pileus, the sequestrate state of the fruit body and the presence of basidiospore ornamentations (see examples in Fig. 1).

**Figure 1.**
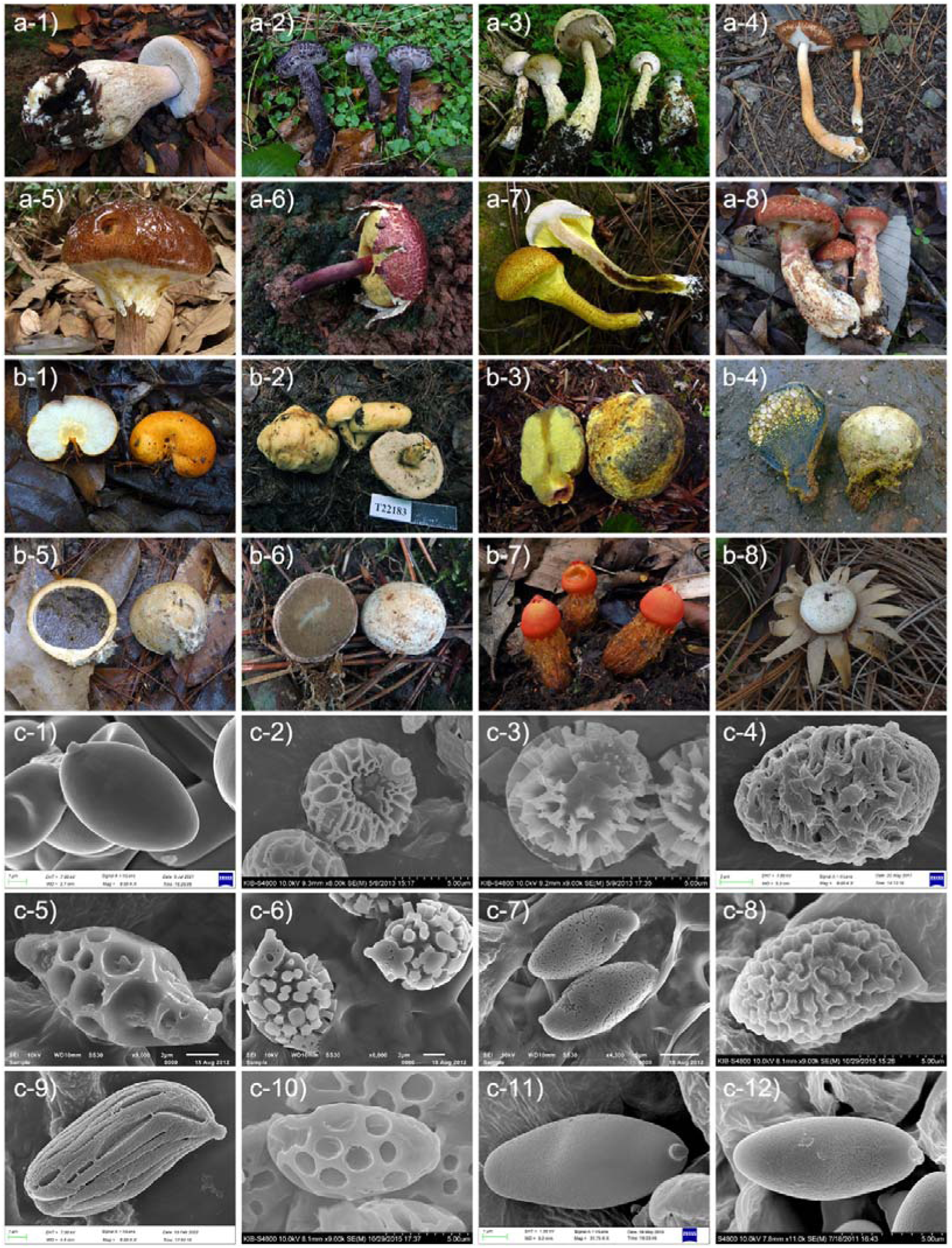
The morphological characters observed in Boletales species. Images a-1 to a-8 depict species without a universal veil (a-1) or with a universal veil (a-2 to a-8). Images b-1 to b-8 showcase species with a sequestrate state of the fruit body. Images c-1 to c-12 present the ornamented basidiospores of various species within Boletales. a-1: *Boletus edulis*, a-2: *Strobilomyces glabriceps*, a-3: *Austroboletus olivaceoglutinosus*, a-4: *Veloporphyrellus pseudovelatus*, a-5: *Aureoboletus viscosus*, a-6: *Boletellus areolatus*, a-7: *Pulveroboletus brunneopunctatus*, a-8 *Suillus phylopictus*; b-1: *Turmalinea chrysocarpa*, b-2: *Leccinellum cremeum*, b-3: *Neoboletus thibetanus*, b-4: *Pisolithus orientalis*, b-5: *Scleroderma yunnanense*, b-6: *Rhizopogon boninensis*, b-7: *Calostoma sinocinnabarinum*, b-8: *Astraeus hygrometricus*; c-1: *Amoenoboletus granulopunctatus*, c-2: *Strobilomyces atrosquamosus*, c-3: *Strobilomyces verruculosus*,c-4: *Spongispora temasekensis*, c-5: *Austroboletus dictyotus*, c-6: *Austroboletus fusisporus*, c-7: *Austroboletus olivaceoglutinosus*, c-8: *Aureoboletus shichianus*, c-9: *Boletellus wenshanensis*, c-10: *Heimioporus retisporus*, c-11: *Hemileccinum rugosum*, c-12: *Xerocomus microcarpoides*.

Specifically, we examined if the origins and maintenance of these traits were related to changes in paleo-temperature and aridity. Ancestral state construction was performed using *rayDISC* and plotted using *plotRECON*, which are implemented in the R package *corHMM* (https://rdrr.io/cran/corHMM/).

## RESULTS

### Phylogenomic analysis reveals a clear phylogenetic framework of Boletales

During the phylogenomic analyses, the D1 dataset consisted of 593373 base pairs (bp) from 1327 gene sequences (translated protein sequences) of 207 Basidiomycota species, including 98 species from 83 genera in 15 families of the Boletales order. The D2 dataset included 125076 bp from 286 gene sequences of the same species. The resultant topologies generated by the D1 and D2 datasets were very similar (Fig. 2, Supporting Information Fig. S1, S2). Based on the phylogenomic framework, the group consisted of the orders Lepidostromatales and Atheliales forming a sister clade to the Boletales, followed by the orders Amylocorticiales. Within the Boletales order, the phylogenetic relationships among families or suborders are clearly delimited.

**Figure 2.**
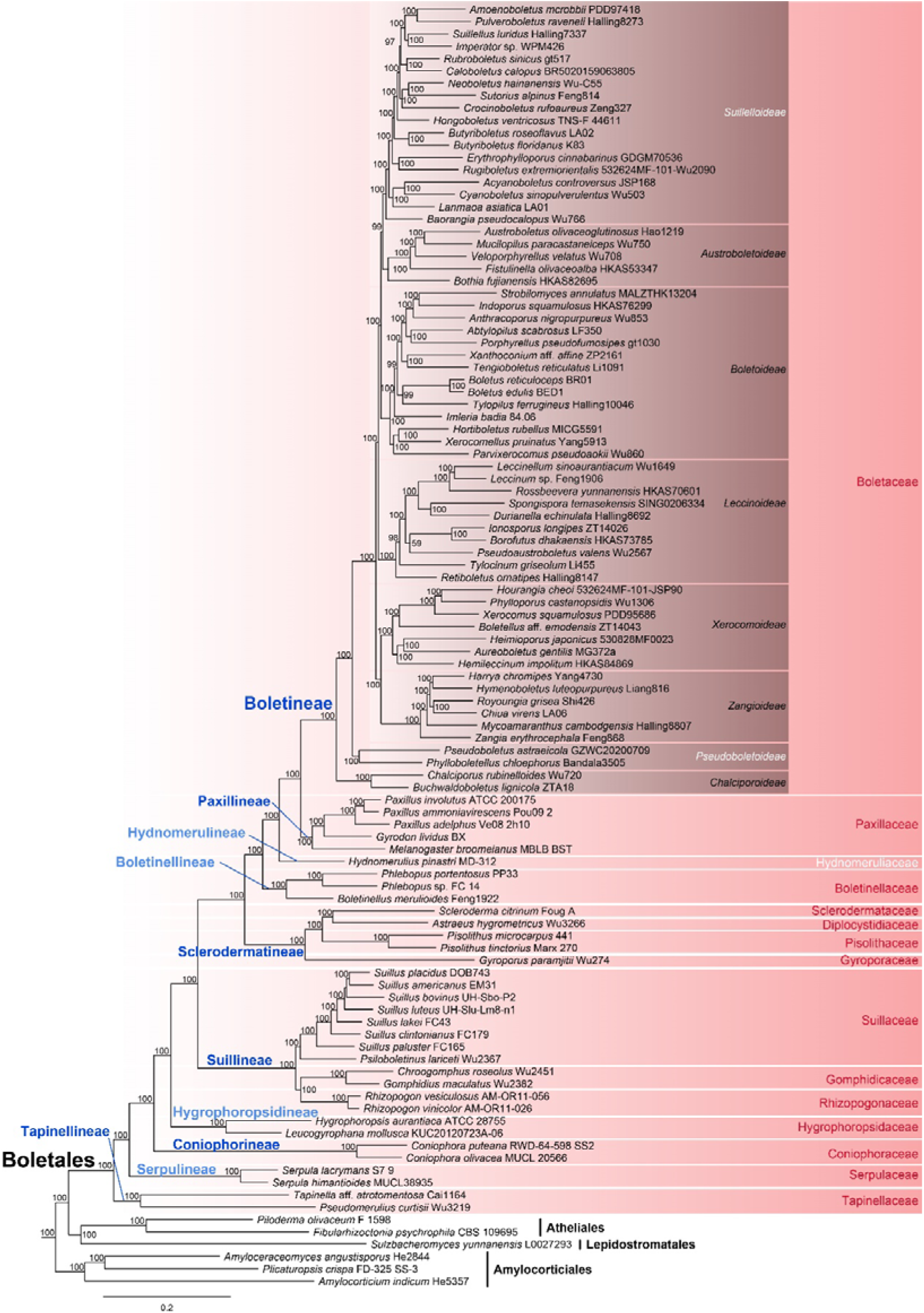
Phylogenomic tree of Boletales using maximum likelihood method based on 1327 gene sequences (a part of Supporting Information Fig. S1). The bootstrap values (>50%) were shown on/below/beside the nodes. The light blue and white letters in the boxes showed the new suborders and new subfamilies, respectively.

Specifically, the suborder Tapinellineae (Tapinellaceae) was uncovered as the most basal group of Boletales, followed successively by clades represented by Serpulineae subord. nov., Coniophorineae (Coniophoraceae), Hygrophoropsidineae subord. nov., Suillineae (including Suillaceae, Rhizopogonaceae and Gomphidiaceae), Sclerodermatineae (including Sclerodermataceae, Diplocystidiaceae, Gyroporaceae), Boletinellineae subord. nov., Hydnomerulineae subord. nov., Paxillineae (Paxillaceae), and finally Boletineae (Boletaceae). Within the family Boletaceae, eight main clades were identified, each receiving strong statistical support. The most basal clade within family Boletaceae was the subfamily *Chalciporoideae*, followed by the *Pseudoboletoideae* subfam. nov. The subfamilies *Zangioideae* and *Xerocomoideae* clustered together and formed the third basal clade. The remaining lineages *Leccinoideae*, *Boletoideae*, *Austroboletoideae* and *Suillelleoideae* subfam. nov. (labelled as the *Pulveroboletus* group in Wu et al. (2014)) were next. Together, utilizing the combined phylogenomic and four-gene data, we generated a highly comprehensive phylogeny of Boletales, including a total of 984 species belonging to 121 genera within 15 families of Boletales (Supporting Information Fig. S3).

### Boletales originates in Early Jurassic with its suborders medially aging in Early Cretaceous

Based on the genome and four-gene dataset, the stem age of order Boletales was similarly estimated to Early Jurassic (185 Mya and 189 Mya, respectively) with a crown age in Middle Jurassic (167 Mya and 163 Mya, respectively). The stem age of suborder Boletineae was also similarly inferred by the two datasets, to late Cretaceous (86 Mya and 77 Mya, respectively). The origin of other suborders including Boletinellineae, Coniophorineae, Hydnomerulineae, Hygrophoropsidineae, Paxillineae, Sclerodermatineae, Serpulineae, Suillineae, and Tapinellineae were estimated to range from 163 Mya to 77 Mya based on the four-gene dataset, encompassing the period from Late Jurassic to Late Cretaceous (Fig. 3, Supporting Information S5, Table S3). The mean stem age of suborders was 106 Mya in Early Cretaceous. Among the suborders, all saprotrophic ones (Coniophorineae, Hygrophoropsidineae, Serpulineae, and Tapinellineae) originated earlier than ectomycorrhizal ones (Boletinellineae, Paxillineae, Sclerodermatineae, and Suillineae) except the saprotrophic Hydnomerulineae (88 Mya). The families within Boletales originated from around 163 to 55 Mya, with a mean stem age of 85 Mya in late Cretaceous. Among them, the family Boletaceae originated at 77 Mya with the crown age of 69 Mya in Late Cretaceous. The genera of Boletales occurred from 142 to 12 Mya, with the mean stem age being 45 Mya in middle Eocene (Figs. 3, Supporting Information S4, Table S3). Within family Boletaceae, the subfamilies had stem ages ranging from Late Cretaceous (69 Mya) to late Eocene (48 Mya), with the mean age being 53 Mya in early Eocene (Figs. 3, Supporting Information S4, Table S3).

**Figure 3.**
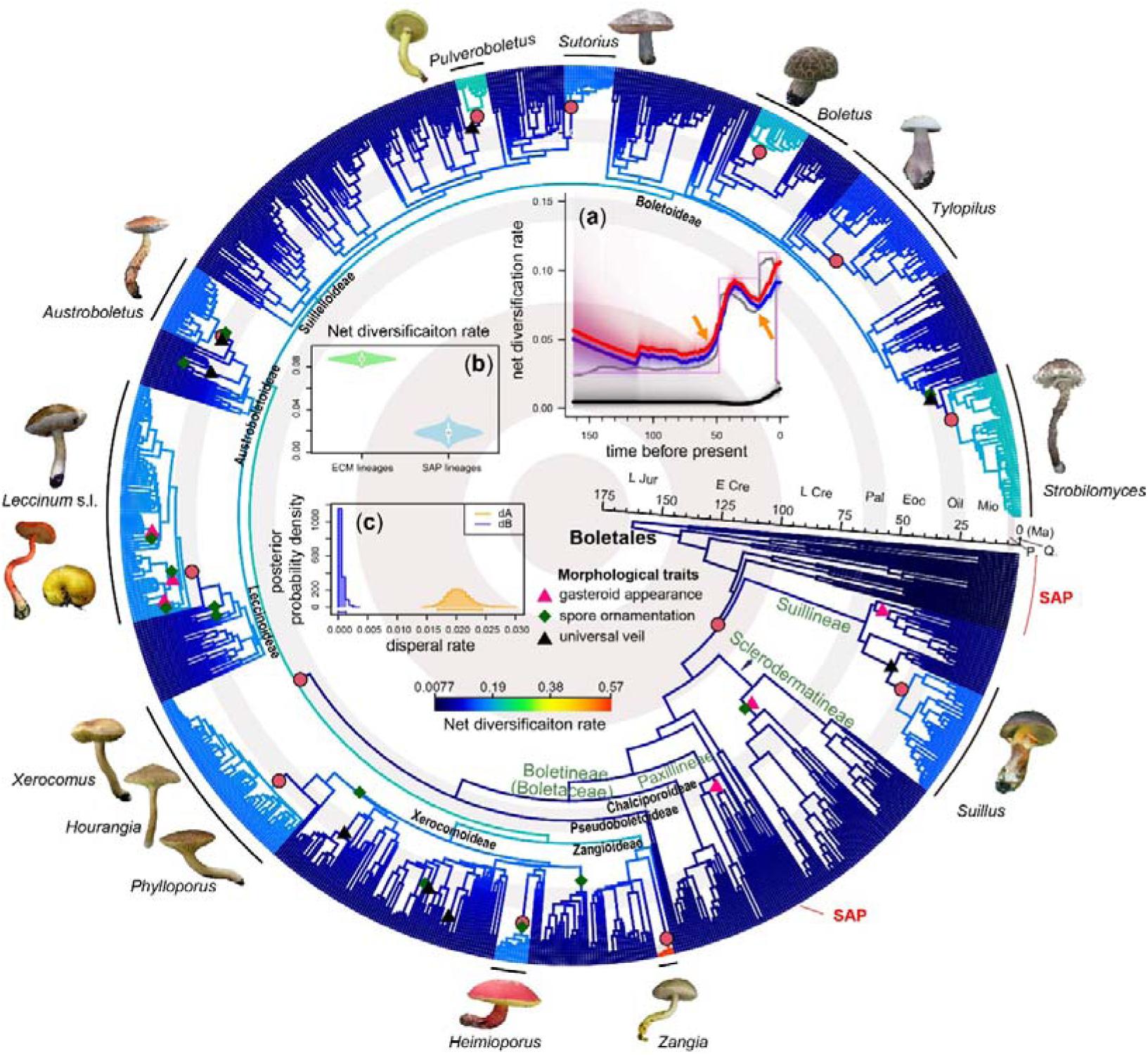
The phylogenetic relationships and diversification patterns across 984 species in Boletales. The tree was constructed using sequences from nrLSU, *TEF1*, *RPB1*, and *RPB2*, with a phylogenomic backbone constraint for higher taxonomic nodes (also refer to Supporting Information Fig. S5). Branches are colored based on the net diversification rate, inferred using Bayesian Analysis of Macroevolutionary Mixtures (BAMM). Warmer colors indicate higher rates of net diversification. Red circles denote significant shifts in diversification rate at specific nodes. The most likely ancestral states of morphological characters, such as sequestrate state of the fruit body (dark pink triangle), ornamented basidiospore (green rhombus), and veiled margin (black triangle), are indicated at the corresponding nodes (also refer to Supporting Information Fig. S10-S12). In the inner part of the plate, *a* shows the diversification curves over geological time for Boletales retrieved from varied methods (red line: speciation rate by *BAMM*, blue line: net diversification rate by *BAMM*, black line: extinction rate by *BAMM*, grey line: net diversification rate by *RevBayes*, lilac line: diversification rate by *TreePar*). Orange arrows indicate the two sharp increases of diversification rate. *b* provides a comparison of net diversification rates between ectomycorrhizal (ECM) and saprotrophic (SAP) lineages. *c* presents the inferred dispersal rates using the GeoSSE model implemented in *diversitree* (orange columns: East Asia, blue columns: the region outside of East Asia). The lineages with a core shift in diversification rate are exhibited at the outermost layer of the plate. In addition, the saprotrophic (SAP) clades are highlighted with red lines at the outermost layer, while the remaining clades represent ectomycorrhizal (ECM) associations.

### ECM lifestyle significantly promoted the lineage diversification and maintenance in Boletales

The rate-through-time plots, generated using three different approaches (*BAMM*, *RevBayes* and *TreePar*), consistently revealed two sharp increases of diversification rate during the periods from ca. 54 Mya to 36 Mya in Eocene and from 17 Mya (early Miocene) to present (Fig. 3a, orange arrows). The estimations of episodic diversification rate, based on the Birth-Death model, using both Bayesian (*RevBayes*) and maximum likelihood (*TreePar*) approaches, revealed an additional significant decline in diversification rate occurring at around 3 Mya, marking the most recent divergence event (Fig. 3a). The diversification pattern of Boletales was mainly driven by increased species number in the ectomycorrhizal suborders Boletineae and Suillineae (Supporting Information Fig. S6), while in Boletineae, the species increases were in subfamilies *Austroboletoideae*, *Boletoideae*, *Leccinoideae, Suillelloideae*, *Xerocomoideae* and *Zangioideae* (Supporting Information Fig. S7). Overall, the global diversification rates of all ectomycorrhizal lineages of Boletales were significantly higher than those of saprotrophic Boletales lineages (Fig. 3b, Supporting Information Fig. S6). The BAMM analysis further identified 13 core diversification shifts, of which all were detected in ectomycorrhizal clades and almost all were in the suborder Boletaceae except one in suborder Suillineae (Fig. 3). Similar core shifts were also uncovered by the estimation of branch-specific diversification rate using the Birth-Death model by RevBayes (Supporting Information Fig. S8). These shifts mainly occurred in three historical periods, namely early Cretaceous, Ecocene and Miocene (Fig. 3). The first shift occurred at around 111 Mya during Early Cretaceous when the ectomycorrhizal mode formed in Boletales, and the second shift appeared at 54 Mya during early Eocene which was dominated by the clade of family Boletaceae. Other shifts aged from late Eocene to Miocene and existed in the generic clades such as *Suillus* in Suillineae, and *Austroboletus*, *Boletus, Hemioporus*, *Leccinum* s.l., *Phylloporus/Hourangia/Xerocomus*, *Pulveroboletus*, *Strobilomyces*, *Sutorius*, *Tylopilus*, and *Zangia* in Boletaceae.

Regarding geographic patterns, the pronounced lineage accumulation in Boletales primarily stems from species in East Asia and North America (Fig. 4, Supporting Information Table S4). Boletales species in East Asia further demonstrated higher rates of speciation and extinction in contrast to the region outside East Asia. The dispersal rate from East Asia to the region outside East Asia surpassed the rate of the opposite dispersal direction (Fig. 3c, Supporting Information Fig. S9).

**Figure 4.**
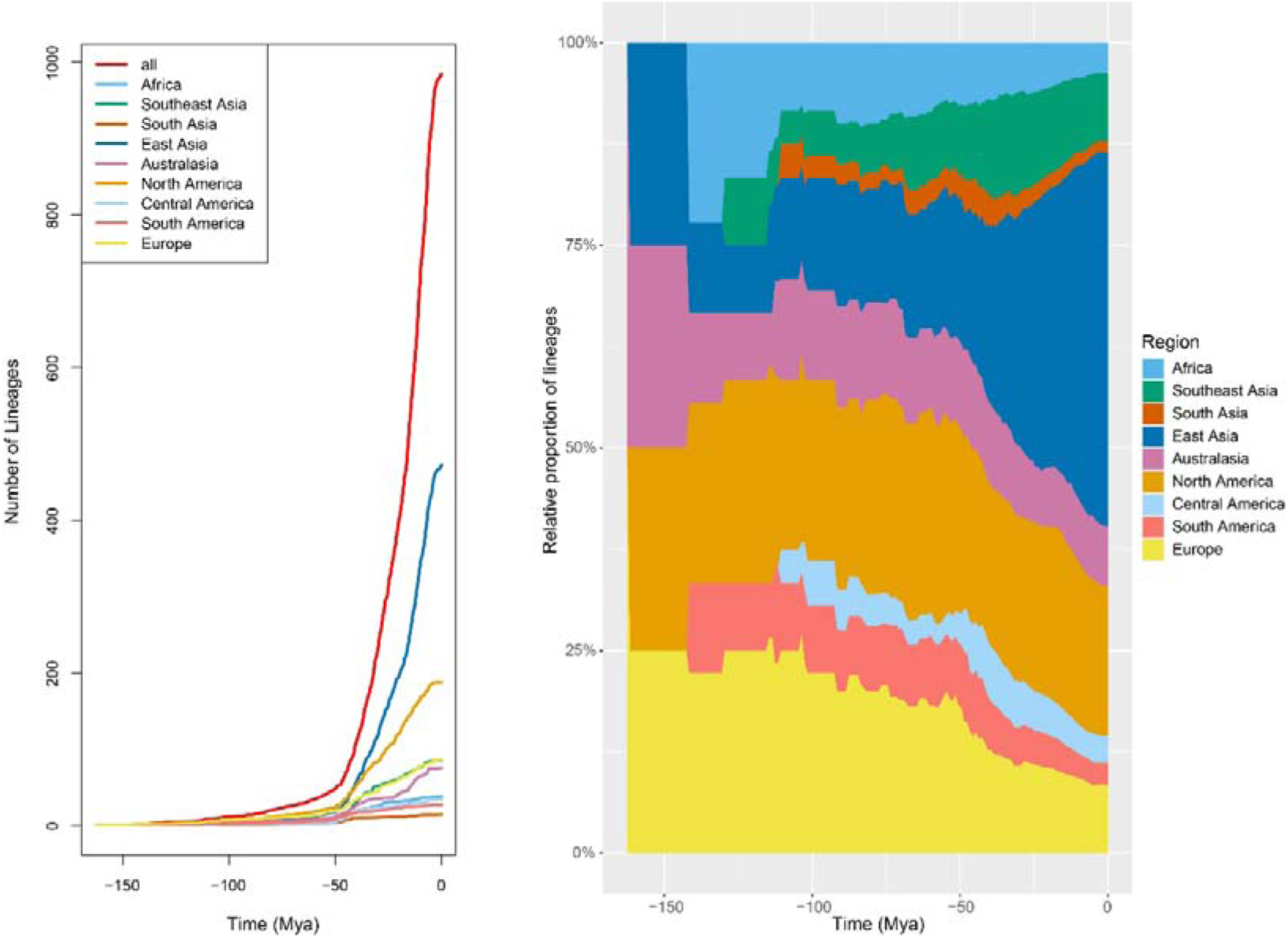
The lineage accumulation of Boletales across different regions over geological time. The left part displays the number of lineages in Boletales across geological time for each region in the world as well as the whole world. The right part illustrates the relative proportion of lineages in Boletales across geological time for each region. The regions included here are Africa, Southeast Asia, South Asia, East Asia, Australasia, North America, Central America, South America, and Europe.

### The rapid diversification of Boletales is correlated with warm-humid paleoclimates and the presence of diverse host plants

Diversification analyses of host plants revealed intriguing patterns in the diversification rates of Fagales. They exhibited a gradual acceleration until the middle Miocene (12 Mya), followed by a significant surge until late Miocene (5 Mya), eventually reaching a plateau phase. Similarly, Pinaceae displayed a progressive diversification acceleration until Early Miocene (23 Mya), followed by a noteworthy increase until the middle Miocene (10 Mya), and subsequently underwent a distinct decline. Employing four distinct environmentally-dependent diversification models (GMRF, HSMRF, UC, and *fixed*) in RevBayes to assess the correlation between diversification rates and potential factors (Fig. 5b), several interesting findings emerged. Firstly, the correlation factor (β) between the diversification rate of Boletales and paleo-temperature was significantly positive (β>0), with a posterior probability (PP) > 0.95 for all models except GMRF, which had a posterior probability of 0.77. Conversely, the correlation between the rate of Boletales and paleo-aridity displayed a moderately to significantly negative relationship (β<0), with a PP ranging from 0.59 to 1. When considering the host plants, there was moderate to significant support for a positive correlation between the rates of Boletales and Pinaceae (β>0, PP=0.65∼1). Similarly, the positive correlation between the rates of Boletales and Fagales tended to be moderate to significant (β>0, PP=0.54∼0.97 for GMRF, HSMRF, and *fixed* models, while PP=0.37 for UC model).

**Figure 5.**
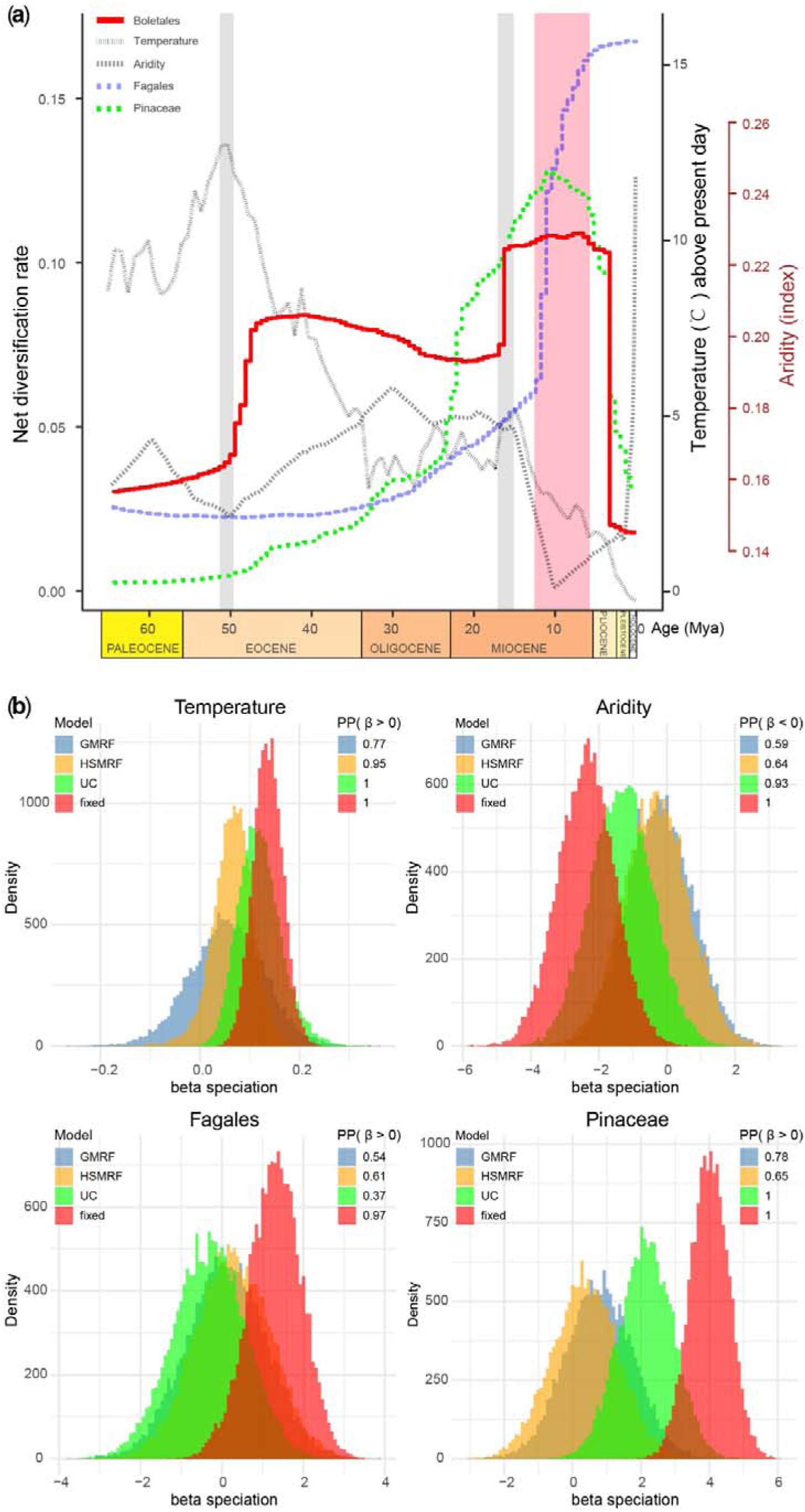
Estimating the diversification rate of Boletales and its correlation to paleo-temperature, paleo-aridity, and the diversification rates of Fagales and Pinaceae by using RevBayes. a) The diversification rates over time for Boletales (red solid line), Fagales (lilac dotted line), Pinaceae (green dotted line), and the fluctuations in paleo-temperature (black dotted line) inferred by Zachos *et al*. (2001) and paleo-aridity (grey dotted line) simulated by Hagen *et al*. (2021). The gray box represents the approximate time of the two sharp increases in diversification during the Eocene and Miocene, while the pinkish box represents the approximate time of sustained highest diversification rates. b) Correlation analyses between the diversification rate of Boletales and four different potential factors (paleo-temperature, paleo-aridity, diversification of Fagales and Pinaceae). Each factor is evaluated using four different models (GMRF, HSMRF, UC, and *fixed*) in separate analyses. The posterior probabilities of the correlation factors (β) (> 0: positively correlated or < 0: negatively correlated) are shown in the sub-panels.

### Key morphological traits of Boletales originated during the Oligocene-Miocene period, with multiple independent evolutionary events

Through analysis and review, it was discovered that the character of the presence of universal veil is predominantly found in the genus *Suillus* of Suillineae, and in the subfamilies of Boletineae, namely *Austroboletoideae*, *Boletoideae*, *Suillelloideae*, and *Xerocomoideae*. The sequestrate state of the fruit body is mainly observed in the genus *Rhizopogon* of Suillineae, most clades of Sclerodermatineae, some clades of Paxillineae and the subfamily *Leccinoideae* of Boletineae. The presence of ornamentations on the surface of basidiospores predominantly exists in most clades of Sclerodermatineae, and the subfamilies of Boletineae, namely *Xerocomoideae*, *Leccinoideae*, *Austroboletoideae*, and *Boletoideae* (Supporting Information Table S6).

Ancestral character reconstruction indicated that the presence of the universal veil, the sequestrate state of the fruit body and the ornamented basidiospores mostly originated during Oligocene to early Miocene. In detailed, the presence of universal veil on pileus independently evolved at least seven times in Boletales and originated between Middle Eocene (39 Mya) and Miocene (12 Mya). Gasteromycetation independently evolved more than five times in Boletales, and originated between Late Cretaceous (74 Mya) and late Miocene (12 Mya). The presence of basidiospore ornamentation independently evolved at least 13 times in Boletales, and originated between Late Cretaceous (74 Mya) and Miocene (11 Mya) (Fig. 3, Supporting Information Fig. S10∼S12). In addition, sporadic cases of gasteromycetation and formation of basidiospore ornamentations also mainly occurred in the Oligocene (Supporting Information Fig. S5, Table S6), such as *Neoboletus thibetanus* (sequestrate state of the fruit body, 23.96 Mya), *Mycoamaranthus cambodgensis* (sequestrate state of the fruit body, 32.95 Mya), *Costatisporus cyanescens* (sequestrate state of the fruit body, 30.12 Mya), *Afrocastellanoa ivoryana* (sequestrate state of the fruit body + ornamented basidiospores, 33.11 Mya), *Amylotrama clelandii* (sequestrate state of the fruit body + ornamented basidiospores, 31.18 Mya), *Heliogaster columellifer* (sequestrate state of the fruit body, 20.09 Mya), *Solioccasus polychromus* (sequestrate state of the fruit body + ornamented basidiospores, 38.21 Mya), and *Phylloboletellus chloephorus* (ornamented basidiospores, 55.32 Mya).

### Frequent reticulate evolution occurred in highly-diversified ectomycorrhizal lineages of Boletales: Boletineae and Suillineae

Based on phylogenetic network analysis using 286 gene trees, it was observed that the ectomycorrhizal (EcM) lineages of Boletales, viz. Boletineae and Suillineae, underwent frequent reticulate evolution during their early evolutionary histories (Fig. 6). Reticulate evolution in Boletineae occurred mostly among the subfamilies *Austroboletoideae*, *Leccinoideae*, *Suillelloideae*, *Xerocomoideae*, and *Zangioideae*, while that in Suillineae occurred among Suillaceae and Rhizopogonaceae.

**Figure 6.**
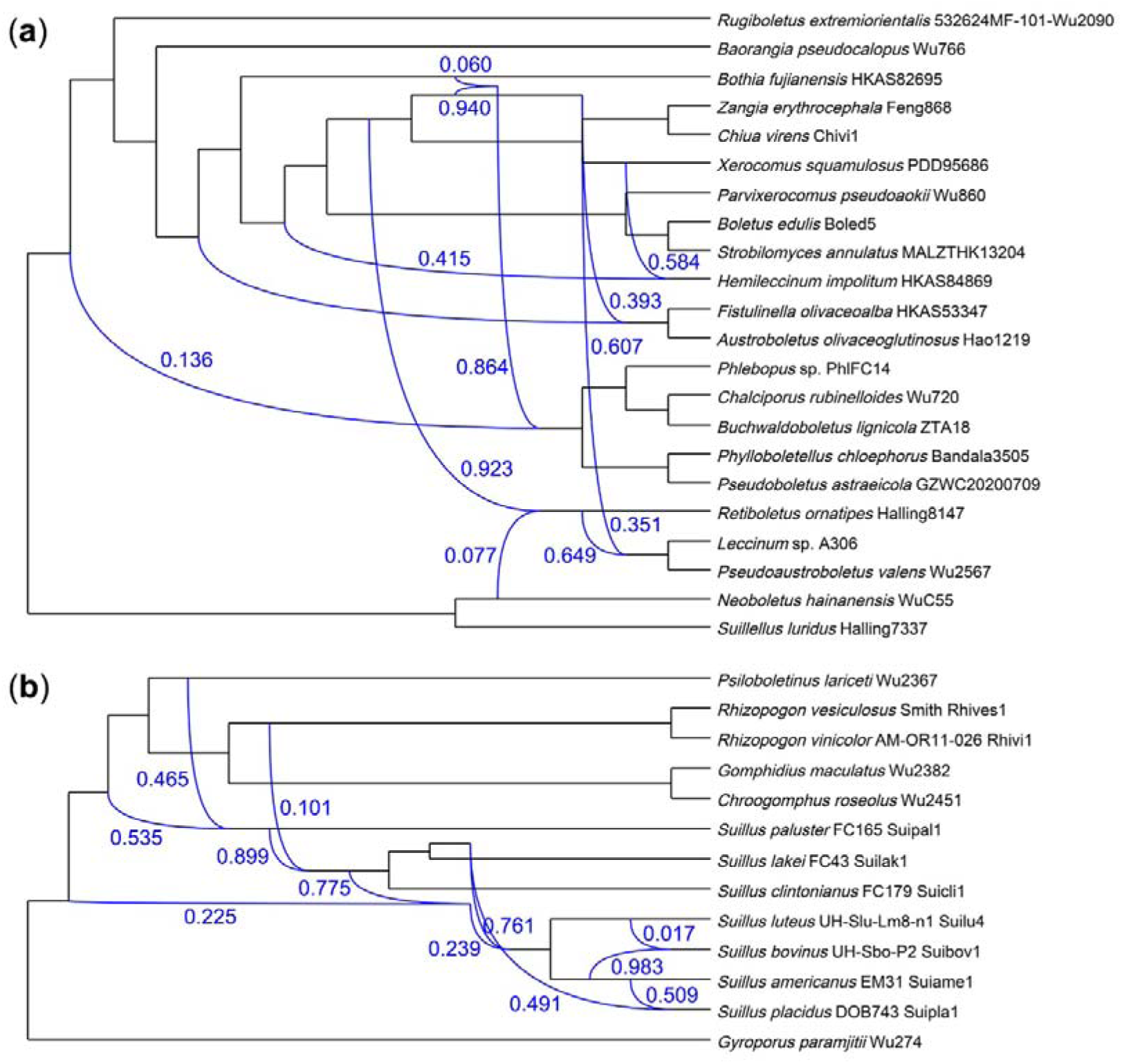
Phylogenetic network analyses of Boletaceae (a) and Suillineae (b). The blue lines represent occurrences of reticulate evolution. The values alongside the blue lines indicate inheritance probabilities.

## TAXONOMY

Based on the findings from our large-scale phylogenomic and phylogenetic analyses, we identified the need to propose four new suborders within the Boletales order.

These suborders are named Boletinellineae, Hydnomerulineae, Hygrophoropsidineae, and Serpulineae. Additionally, we erected a new family called Hydnomeruliaceae, which belongs to the Hydnomerulineae suborder, and two new subfamilies, *Pseudoboletoideae* and *Suillelloideae*, within the Boletaceae family of the Boletineae suborder. These taxonomic treatments were based on an integration of molecular and morphological evidence and help accommodate previously unclassified lineages.

**Boletinellineae** G. Wu & Zhu L. Yang, subord. nov.

*MycoBank ID*: 849907

*Typus*: *Boletinellus* Murrill, Mycologia 1(1): 7 (1909)

*Description*: Basidioma boletoid to boletinoid. Pileus smooth or subtomentose, dry; marginal veil absent or margin incurved; context yellowish to yellow, unchanged or bluing when bruised; hymenophore tubular, boletoid, or boletinoid with radial arrangement, surface yellowish, yellow or brown, tube yellowish to yellow, unchanged or bluing when bruised. Stipe stout or slim, without reticulations.

Pileipellis trichodermium. Basidiospores broadly elliptical, yellowish brown to light brown in 5% KOH. Hymenophoral trama generally boletoid. Clamp connections present. On soil.

*Exemplar family*: Boletinellaceae P.M. Kirk *et al*.

*Exemplar genera*: *Boletinellus* Murrill, *Phlebopus* (R. Heim) Singer

**Hydnomerulineae** G. Wu & Zhu L. Yang, subord. nov.

*MycoBank ID*: 849908

*Typus*: *Hydnomerulius* Jarosch & Besl, Pl. Biol. 3(4): 447 (2001)

*Description*: Basidioma resupinate, effused. Hymenophore surface smooth to hydnoid, yellow to brownish yellow, brownish staining when injured. Basidiospores cyanophilous and weakly dextrinoid, thin-walled. Sclerotia present. Hyphal strands present. Clamp connections common. On rotten wood. Brown-rot.

*Exemplar family*: Hydnomeruliaceae G. Wu & Zhu L. Yang

*Exemplar genus*: *Hydnomerulius* Jarosch & Besl

**Hygrophoropsidineae** G. Wu & Zhu L. Yang, subord. nov.

*MycoBank ID*: 849909

*Typus*: *Hygrophoropsis* (J. Schröt.) Maire ex Martin-Sans, L’Empoisonnem. Champ.: 99 (1929)

*Description*: Basidioma pileate-stipitate or resupinate. When pileate-stipitate, pileus applanate, usually depressed in the center, often incurved at margin, dry, subtomentose, whitish, cream to yellowish, yellow, ochre-yellow. Context whitish, unchanged when hurt. Hymenophore lamellate, forked, decurrent, unchanged when hurt. Stipe subcylindrical, scaly or smooth. Basidiospores ellipsoid, smooth, dextrinoid. Pileipellis irregularly interwoven. Clamp connections common. On soil. When resupinate, basidioma yellow-orange or orange-brown. Hymenophore merulioid, surface smooth. Basidiospore ellipsoid, cyanophilous, dextrinoid. Clamp connections common. On rotten wood.

*Exemplar family*: Hygrophoropsidaceae Kühner

*Exemplar genera*: *Hygrophoropsis* (J. Schröt.) Maire ex Martin-Sans,

*Leucogyrophana* Pouzar

**Serpulineae** G. Wu & Zhu L. Yang, subord. nov.

*MycoBank ID*: 849910

*Typus*: *Serpula* (Pers.) Gray, Nat. Arr. Brit. Pl. (London) 1: 637 (1821)

*Description*: Basidioma resupinate, effused-reflexed, up to pileate. Hymenophore meruliod to poroid, or tuberculate, yellowish brown to brown. Basidiospores broadly elliptical to slight ovoid, yellowish, cyanophilous, thick-walled. Clamp connections common. On rotten wood. Brown-rot.

*Exemplar family*: Serpulaceae Jarosch & Bresinsky

*Exemplar genus*: *Serpula* (Pers.) Gray

**Hydnomeruliaceae** G. Wu & Zhu L. Yang, fam. nov.

*MycoBank ID*: 849911

*Typus*: *Hydnomerulius* Jarosch & Besl, Pl. Biol. 3(4): 447 (2001)

Description: Basidioma resupinate, effused. Hymenophore surface smooth when young, becoming hydnoid with age, yellow to brownish yellow, brownish staining when injured. Basidiospores broadly elliptical, pale yellow in 5% KOH, cyanophilous and weakly dextrinoid, thin-walled. Sclerotia numerous, typically ovoid to broadly ellipsoid. Hyphal strands present, brown to blackish. Clamp connections common. On rotten wood. Brown-rot.

*Exemplar genus*: *Hydnomerulius* Jarosch & Besl

***Pseudoboletoideae*** G. Wu, Halling & Zhu L. Yang, subfam. nov.

*MycoBank ID*: 849912

*Typus*: *Pseudoboletus* Šutara, Česká Mykol. 45(1-2): 2 (1991)

*Description*: Basidioma boletoid or phylloporoid. Pileus viscid when wet, smooth, or subtometose; marginal veil absent, sometimes margin incurved; context whitish, yellowish to yellow, staining blue or unchanged when cut; hymenophore tubular or lamellate, yellowish to yellow, staining blue or unchanged when cut. Stipe subcylindrical, scaly. Pileipellis ixotrichodermium to ixocutis. Basidiospores short ellipsoid to fusoid, smooth or longitudinally striate, yellowish brown in 5% KOH. Hymenophoral trama boletoid. On soil. fungal-parasite or non-ectomycorrhizal.

*Exemplar genera*: *Phylloboletellus* Singer, *Pseudoboletus* Šutara

***Suillelloideae*** G. Wu, Halling & Zhu L. Yang, subfam. nov.

*MycoBank ID*: 849913

*Typus*: *Suillellus* Murrill, Mycologia 1(1): 16 (1909)

*Description*: Basidioma boletoid, or rarely gasteroid. Pileus dry, smooth, subtometose, or pulverulent; marginal veil usually absent, sometimes present, or margin incurved. Context often yellowish to yellow, occasionally white to grayish, usually staining blue when cut, occasionally unchanged. Hymenophore tubular or rarely glebulous, tube yellowish to yellow, occasionally whitish to grayish or pinkish, pores yellowish to yellow, sometimes brownish, brown, or reddish brown, occasionally vinaceous brown to grayish brown, often staining blue when cut, occasionally unchanged. Stipe smooth, reticulate, or pulverulent. Pileipellis generally trichodermium, sometimes subcutis, rarely ixotrichodermium. Basidiospores often subfusoid, or globose to subglobose, ellipsoid to broadly ellipsoid, usually smooth, yellowish brown in KOH. Hymenophoral trama boletoid or phylloporoid. When gasteroid, basidioma often with a rudimentary stipe or rhizomorph. Gleba yellowish to yellow, or brown to violet brown, bluing when cut. Basidiospores smooth or longitudinally striate. Clamp connections absent. On soil. Ectomycorrhizal.

*Exemplar genera*: *Acyanoboletus* G. Wu & Zhu L. Yang, *Amoenoboletus* G. Wu, *et al*., *Baorangia* G. Wu & Zhu L. Yang, *Butyriboletus* D. Arora & J.L. Frank, *Cacaoporus* Raspé & Vadthanarat, *Caloboletus* Vizzini, *Costatisporus* T.W. Henkel & M.E. Sm., *Crocinoboletus* N.K. Zeng, *et al*., *Cyanoboletus* Gelardi, *et al*., *Erythrophylloporus* Ming Zhang & T.H. Li, *Gymnogaster* J.W. Cribb, *Hongoboletus* G. Wu & Zhu L. Yang, *Imperator* Koller *et al*., *Lanmaoa* G. Wu & Zhu L. Yang, *Neoboletus* Gelardi, *et al*., *Pulveroboletus* Murrill, *Rubroboletus* Kuan Zhao & Zhu L. Yang, *Rugiboletus* G. Wu & Zhu L. Yang, *Singerocomus* T.W. Henkel & M.E. Sm., *Suillellus* Murrill, *Sutorius* Halling, *et al*.

## DISCUSSION

The order Boletales is a globally recognized group of fungi that is famous for its ecological importance, high-value edibility, frequent poisoning, and complex classification (Nuhn *et al*., 2013; Wu *et al*., 2014). An 87-single-copy-gene phylogenetic tree of Boletales was constructed previously (Sato & Toju, 2019), and recently the first phylogenomic tree specific to Boletales was inferred using 28 whole genomes from 27 different species (Wu *et al*., 2022). However, the evolutionary relationships within the highly diversified Boletaceae family are not yet fully resolved. In this study, we reconstructed a high-resolution phylogenomic tree of Boletales using deep genome-skimming data with a comprehensive sampling at the generic level.

Consistent with the findings of Wu *et al*. (2022), all known brown-rot fungi of Boletales occupy the basal lineages of the phylogenetic tree, with the exception of *Hydnomerulius pinastri*, which is clustered among ectomycorrhizal lineages. In the Boletaceae family, our results confirm the monophyly of the seven clades at subfamily level proposed by Wu *et al*. (2014) and further clarify the relationships among these main clades, with the inclusion of a newly identified subfamily called *Pseudoboletoideae*. In addition, we determine that the Asian parasite bolete *Pseudoboletus astraeicola* and Mexican *Phylloboletellus chloephorus* cluster closely with *Pseudoboletus parasiticus* for the first time, providing further evidence for the partial parasitic ability of the basal lineages of Boletaceae (Nuhn *et al*., 2013). Using genome-skimming data, we finally confirm with robust support that the *Pulveroboletus* group, as previously circumscribed by Wu *et al*. (2014), is a monophyletic clade and in the present work formally name it *Suillelloideae*. This study also verified that the unclassified lineage consisting of *Bothia*, *Phylloporopsis*, and *Solioccasus* unambiguously clustered within the subfamily *Austroboletoideae*.

With these new findings, we have robustly elucidated the phylogenetic relationships among the eight subfamilies of Boletaceae and those among the 10 suborders and 15 families of Boletales.

Compared to the divergence times estimated by Varga *et al*. (2019) and Sánchez-García *et al*. (2020) using genome-scale data, the mean stem age of Boletales (185 Mya) estimated in this study is slightly older than theirs, 21 Mya older than 164 Mya of Varga et al. (2019) and 14 Mya older than 171 Mya of Sánchez-García et al. (2020). In the evolutionary history of Boletales, the previous study (Sato & Toju, 2019) has suggested that there was a rapid diversification of Boletaceae. Our analysis confirms it and further reveals that this process was not uniform but underwent two distinct episodes which began at Early Eocene (ca. 54 Mya) and middle Miocene (ca. 17 Mya), respectively (Fig. 3a). Interestingly, these two periods coincide with two well-known climatic optima: the Early Eocene Climatic Optimum (EECO, approximately 53–50 Mya) (Zachos *et al*., 2001; West *et al*., 2020) and the Mid-Miocene Climatic Optimum (MMCO, approximately 17–14 Mya) (Zachos *et al*., 2001; Böhme, 2003). Both of these warming events created relatively warm and humid environments, which had a significant impact on the evolution of global biodiversity. During the EECO, for example, Theaceae plants (Cheng *et al*., 2022) and insects (Wang *et al*., 2014) underwent early radiation, and primates reached a peak in species diversity during the MMCO (Merceron *et al*., 2012). It is worth noting that the host plants of Boletales–Fagales and Pinaceae–also experienced radiation during the MMCO (Jin *et al*., 2021; Zhou *et al*., 2022; This study). Additionally, the MMCO is believed to have strongly influenced the East Asian biota through the northern expansion of the megathermal rainforest biome (Wang *et al*., 2021).On the other hand, the complex tectonic activity during the Eocene and Miocene, such as the uplift of Tibet-Himalaya-Hengduan region, Andes, Western North American Rockies & Sierras, European Alpine System, and the collision between Australia and SE Asia, increased habitat heterogeneity, which contributed significantly to species radiation of plants and animals (Vargas, 2003; Hall, 2011; Hughes & Atchison, 2015; Ding *et al*., 2020; Ding *et al*., 2022). These two widely recognized climatic optima, along with the heterogeneity in geography and plant distributions, could collaboratively impact the diversification of fungi.

In our analysis, it is inferred that the region of East Asia plays a significant role in the diversification of Boletales (Fig. 3c, Supporting Information Fig. S9), which was also noted in previous studies on biogeography of several boletoid fungal genera, such as *Strobilomyces* (Han *et al*., 2018) and *Boletus* (Feng *et al*., 2012). Specifically, we showed that the mean modeled annual precipitation during Early Cretaceous to the present in East Asia (Farnsworth *et al*., 2019) largely coincided with the diversification pattern of Boletales (Fig. 3a). In addition, the intensification of the Asian monsoon from the middle-Miocene to early Pliocene (15 Mya-3Mya) (Sun & Wang, 2005) overlapped with the second rapid diversification event within Boletales (Fig. 3a). Therefore, paleo-precipitation can be an important driver of the diversification of Boletales.

The evolution of mycorrhizal fungi closely associated with host plants is likely influenced by the evolution of their hosts (Sánchez-Ramírez *et al*., 2015; Sánchez-García & Matheny, 2017; Wilson *et al*., 2017). Although pileate-stipitate form of fungi is considered as a key innovative character to promote the diversification of the Agaricomycetes in general (Varga *et al*., 2019; Sánchez-García *et al*., 2020), host-shift and rapid host diversification have shown to play important roles on the diversification of several groups of fungi, such as Boletaceae, *Entoloma*, *Laccaria* and *Tricholoma* (Sánchez-García & Matheny, 2017; Wilson *et al*., 2017; Sato & Toju, 2019; Varga *et al*., 2019) including the whole class Agaricomycetes (Sato 2023). During the late Eocene and late Miocene, especially the latter episode, Fagales, a host plant group of Boletales, diversified rapidly, particularly the lineages within families Betulaceae (Yang *et al*., 2019; Yang *et al*., 2022) and Fagaceae (Cavender-Bares *et al*., 2015; Deng *et al*., 2018; Yang *et al*., 2018; Zhou *et al*., 2022). *Pinus*, another host plant group, also diversified rapidly, with nearly 90% of its species originating in the Miocene era (Jin *et al*., 2021). Because of the close association between mycorrhizal fungi and plants (Brundrett, 2002) and the significant correlations observed here between the diversification rates of Boletales and the above two groups of plants (Fig. 5b), we believe that the accelerated diversification of host plants facilitated the speciation of ectomycorrhizal Boletales.

Environmental change exerts significant influence on the diversification of organisms (Folk *et al*., 2019). In parallel, organisms accordingly need to evolve genetic mechanisms to adapt in the diversified ecological niches. Reticulate evolution emerges as one of the notable adaptive strategies, specifically through the process of introgressive hybridization (Rocha *et al*., 2023). By incorporating genetic material from different species, reticulate evolution accelerates the formation of novel traits and promotes the expansion of biodiversity. This process has been documented in various taxa, including *Diplostephium* (Vargas *et al*., 2017), Fagaceae (Zhou *et al*., 2022), *Picea* (Sun *et al*., 2018), Conidae (Wood & Duda, 2021) and foxes (Rocha *et al*., 2023). In our study, Boletineae and Suillineae, the most diverse groups within Boletales, have also experienced prominent reticulate evolution (Fig. 6), underscoring the importance of this process in the rapid diversification of fungi.

The Boletaceae is renowned for its taxonomic complexity, primarily due to its diverse morphologies and frequent occurrence of convergent evolution (Wu *et al*., 2014). Several morphological characteristics, such as a universal veil of the pileus, sequestrate state of the fruiting body, and ornamentation of basidiospores, likely contributed to the taxonomic complexity and accordingly received significant attention from mycologists. Our analyses suggest that these three traits mainly originated in the arid period from Oligocene to Early Miocene (Fig. 5a). In addition, we observed that the sequestrate state of the fruit body and the presence of a universal veil frequently co-occur with the formation of basidiospore ornamentations during the evolutionary history of Boletales (Fig. 2, Supporting Information S10-S12, Table S6). For example, *Astraeus*, *Calostoma*, *Chamonixia*, *Durianella*, *Octaviania*, *Pisolithus*, *Rhodactina*, *Rossbeevera*, *Scleroderma*, *Spongiforma*, *Tremellogaster* and *Turmalinea* all exhibit a sequestrate state of the fruit body and ornamented basidiospores, while *Austroboletus*, *Boletellus*, and *Strobilomyces* have a universal veil and basidiospore ornamentation. Therefore, we can deduce that these three crucial morphological characteristics are adaptations that Boletales species have developed throughout their evolutionary history in response to arid climates. Specifically, we hypothesize that these three traits allowed the fruiting bodies to maintain moisture and enable the development of mature basidiospores.

In this study, while acknowledging the inherent limitations of our sampling efforts, we deliberately downplayed the significance of geographic distribution in assessing Boletales species. However, we recognize that exploring the geographical dimension is crucial in unraveling the intricacies of their diversification. With a future endeavor of more comprehensive sampling, a fascinating tapestry of regional diversification patterns within the Boletales order awaits our discovery. Furthermore, by merging intricate details pertaining to regional host plants, historical temperature fluctuations, and precipitation patterns, we hold the potential to illuminate a more resplendent panorama of the diversification drivers shaping the remarkable evolution of Boletales.

## DATA AVAILABILITY

The sequences underlying this article are available in GenBank Nucleotide Database (see Supporting Information Table S1 and S2). The phylogenetic trees are available in TreeBase (XXXX).

## ACKNOWLEDGEMENTS

This work was supported by Yunnan Xingdian Talents Support Plan - Science and Technology Leading Talents Program (grant number 202305AB350004) (to Z.L.Y.), the National Natural Science Foundation of China (grant number 31970015, 32270025) (to G.W.), the Yunnan Ten Thousand Talents Program Plan Young & Elite Talents project (YNWR-QNBJ-2018-266) (to G.W.), the Natural Science Foundation of Yunnan Province (202301AW070011, 202201AT070128) (to G.W.), CAS “Light of West China” Program (to G.W.). The authors thank Drs. Ting Guo, Yan-Jia Hao, Pan-Meng Wang, Nian-Kai Zeng at Kunming Institute of Botany, Chinese Academy of Sciences, Dr. Ping Zhang at Hunan Normal University, Dr. Fang Li at Sun Yat-Sen University, Dr. Arooj Naseer at University of the Punjab in Pakistan for collecting samples, and the herbaria GDGM, HICPC, MICG, NY, PDD, ROHB, SING, TNS and ZT for loaning specimens, and the platform for Plant Multi-dimensional Imaging and Diversity Analysis at Kunming Institute of Botany, Chinese Academy of Sciences for scanning basidiospores. The authors are also grateful to the editor and anonymous reviewers for their constructive suggestions.

## AUTHOR CONTRIBUTIONS

Z.L.Y. and G.W. designed the project. G.W. did the analyses. G.W. and Z.L.Y. wrote the manuscript with the help of J.X. and R.E.H.. G.W. coordinated genome sequencing at KIB. K.W., X. X., G.M. L. performed PCR experiments for Sanger sequencing. R.E.H., E.H., S.L., L.P., R.F.A., S.T.N.E., S.A., A.M.P., N.S.Y., B.F., Y.C.L. collected or loaned the important samples for analyses.

## SUPPORTING INFORMATION

Additional Supporting Information may be found online in the Supporting Information section at the end of the article.

**Figure S1.** The phylogenomic tree of Basidiomycota constructed using the maximum likelihood method with 80% (1327) shared genes. Special emphasis was placed on the order Boletales, highlighted within a colored box. Bootstrap values (≥50%) were displayed on/below/beside the nodes. The light blue and white letters within the boxes represented the new suborders and new subfamilies/family, respectively.

**Figure S2.** The phylogenomic tree of Basidiomycota constructed using maximum likelihood method with 90% (286) shared genes. The bootstrap values (≥50%) were displayed on/below/beside the nodes.

**Figure S3.** The phylogenetic tree of Boletales constructed using maximum likelihood method with nrLSU, *TEF1*, *RPB1* and *RPB2* sequences. The bootstrap values (≥50%) were displayed on/below/beside the nodes.

**Figure S4.** The calibrated phylogenomic tree of Boletales constructed using MCMCtree with 90% shared genes. Nodes ages (Mya) were displayed next to the nodes, with blue node bars representing 95% confidence intervals.

**Figure S5.** The calibrated phylogenetic tree of Boletales constructed using RelTime-ML method of MEGA 11 with nrLSU, *TEF1*, *RPB1* and *RPB2* sequences. Nodes ages (Mya) were displayed next to the nodes, with blue node bars representing 95% confidence intervals.

**Figure S6.** Estimated branch-specific diversification pattern across 984 species in Boletales. The tree was constructed using sequences from nrLSU, *TEF1*, *RPB1*, and *RPB2*, with a phylogenomic backbone constraint for higher taxonomic nodes (also refer to Supporting Information Fig. S5). Branches are colored based on the net diversification rate, inferred using RevBayes. Warmer colors indicate higher rates of net diversification. The clades with a significant shift in diversification rate are displayed on the right side of the phylogenetic tree. The Boletaceae family and the ectomycorrhizal lineage of Boletales are marked with black arrows.

**Figure S7.** The diversification patterns of Boletales, highlighting the different suborders within Boletales as well as the evolutionary trajectories of ectomycorrhizal and saprotrophic lineages over geological time. Each plot represents the speciation rate (red line), net diversification (blue line), and extinction rate (black line) estimated through geologic time using the BAMM model and a four-gene dataset including 984 species. The shaded areas indicate the 95% confidence intervals of the rate estimates. The bottom plot specifically compares the net diversification rates among the different suborders within Boletales. It is of note that the phylogenetic positions of the *Austropaxillus* clade and *Penttilamyces* clade within Boletales could not be confirmed by genome data in this study, and thus they were labeled using their generic names.

**Figure S8.** The patterns of diversification within different subfamilies of Boletaceae over geological time. Each plot represents the speciation rate (red line), net diversification (blue line), and extinction rate (black line) estimated using the BAMM model on a four-gene dataset consisting of 814 species. The shaded areas indicate the 95% confidence intervals of the rate estimates. The bottom plot specifically compares the net diversification rates among the different subfamilies within Boletaceae.

**Figure S9.** The results of Geographic State Speciation and Extinction (GeoSSE) modeling for Boletaceae. The upper plot compares the speciation rates between the regions of East Asia and regions outside of East Asia. The middle plot presents the extinction rates, while the bottom plot displays the net diversification rates. The orange columns represent the region of East Asia, while the blue columns represent regions outside of East Asia.

**Figure S10.** The ancestral state reconstruction of the universal veil of pileus in Boletales. The blue pie chart represents the pileus with a universal veil, while the red pie chart represents the pileus without a universal veil.

**Figure S11.** The ancestral state reconstruction of the gasteromycetation (mode of development) of the fruit body in Boletales. The blue pie chart represents the sequestrate state, indicating a closed or partially enclosed fruit body, while the red pie chart represents the pileate-stipitate state, indicating an open and stalked fruit body.

**Figure S12.** The ancestral state reconstruction of basidiospore ornamentation in Boletales. The blue pie chart represents basidiospores with ornamentation, while the red pie chart represents basidiospores without ornamentation (smooth).

**Table S1.** The taxonomy, locality, and assembly information of samples for phylogenomic analysis of Boletales

**Table S2.** The detailed information of samples for four-gene phylogenetic analysis of Boletales. Samples in Boldface are newly sequenced.

**Table S3.** The divergent time of taxonomic clades above generic level of Boletales.

**Table S4.** The proportion of species number in different regions through geological time.

**Table S5.** The inferred mean values of different variants through the geological time from 65 Mya to present used for correlation and influence analyses.

**Table S6.** The matrix for ancestral character reconstruction of key morphologies of Boletales

