## Supplementary material for "The rapid diversification of Boletales is linked to Early Eocene and Mid-Miocene Climatic Optima": Supporting information-Methods S1-plain_re.docx

**New Phytologist Supporting Information**

**Methods S1**

**DNA extraction, sanger sequencing, next-generation-sequencing library preparation and data generation**

For four-gene phylogenetic analysis, total genomic DNA were extracted from dried materials using CTAB method (Doyle & Doyle, 1987). Fragments of four nuclear loci, including nuclear ribosomal large subunit (nrLSU), translation elongation factor 1-α (*TEF1*), RNA polymerase II largest subunit (*RPB1*) and RNA polymerase II second largest subunit (*RPB2*) were amplified using LR0R/LR5, EF1-B-F1 (or EF1-B-F2)/EF1-B-R, RPB1-B-F/RPB1-B-R, RPB2-B-F1 (or RPB2-B-F2)/RPB2-B-R, respectively (Vilgalys & Hester, 1990; Wu *et al.*, 2014). PCR procedures and sequencing for these loci followed the protocols described by Wu et al. (2014). The newly generated sequences were submitted to GenBank (Support Information Table S1, S2).

To perform deep genome skimming, we utilized Next-Generation-Sequencing. Total genomic DNA was extracted from dried materials using either the TIANGEN DNAsecure Plant Kit (DP320, TIANGEN Biotech (Beijing) Co., Ltd.) or the CTAB method (Doyle & Doyle, 1987). All samples were subsequently treated with the protocols described in the NEBNext Ultra™ II DNA library Prep kit for Illumina (NEB #E7645S/L, New England Biolabs) with some minor optimizations following Zeng *et al*. (2018) to create blunt-end DNA libraries in laboratory settings. A total of 64 samples were pooled into dozens of libraries, and paired-end (150 bp) sequencing was performed on one lane for each library using the DNBSEQ-T7 platform (BGI, Shenzhen, China) at the Germplasm Bank of Wild Species, Kunming Institute of Botany, Chinese Academy of Sciences. This generated approximately 3 Gb of data per sample.

**Sequence alignment**

Both the nucleotide sequences of four genes (nrLSU, *TEF1*, *RPB1*, and *RPB2*) and the protein sequences retrieved from genome skimming data were aligned using *mafft* 7.508 with modified settings as follows: --maxiterate 100, --genafpair --reorder (Katoh & Standley, 2013). Poorly aligned bases were trimmed by *trimal* with modified settings as follows: -*automated1*, -*keepseqs* (Capella-Gutiérrez *et al.*, 2009). The cleaned-aligned nucleotide or protein sequences were then separately concatenated using the Perl script *catfasta2phyml.pl* (https://github.com/nylander/catfasta2phyml).

**Dispersal analyses**

Since our sampling could still be limited for a comprehensive biogeographical analysis, we instead focused on studying dispersal events between two regions. One of these regions is East Asia, which is inferred as the center of existing species diversity of boletes, while the other region is outside of East Asia (the rest of world). This approach allows us to gain insights into the respective contributions of these two regions to the diversification of Boletales. We used the *GeoSSE* model (Geographic State Speciation and Extinction) implemented in *Diverstree* v0.9-16 (FitzJohn, 2012) for this analysis, setting the sampling fractions of East Asia and region outside of East Asia to 0.60 (425/704) and 0.29 (389/1328), respectively. These fractions were calculated using the method of Sato and Toju (2019) as described above. The number of MCMC steps was set to 10000.

**Phylogenetic network analyses**

To detect ancient hybridization and introgression, we conducted separate analyses of the phylogenetic networks within the Suillineae and Boletaceae (Boletineae) clades. These two clades encompass the major ECM Boletales species. We employed *PhyloNet* (Solís-Lemus *et al.*, 2017) with a maximum of 6 reticulations and maximum pseudo-likelihood to perform these analyses. For sampling, all 12 Suillineae species in the phylogenomic tree were included with one additional outgroup species, and 21 Boletineae species were selected with one outgroup species such that all subfamilies of Boletaceae were covered and two to four representative genera were chosen in each subfamily. The gene trees used in phylogenetic network analysis were retrieved from gene sequences of D2 dataset.
