## Supplementary figures and images for "The rapid diversification of Boletales is linked to Early Eocene and Mid-Miocene Climatic Optima"

### Fig. S1. Phylogenomic_tree_0.80.jpg

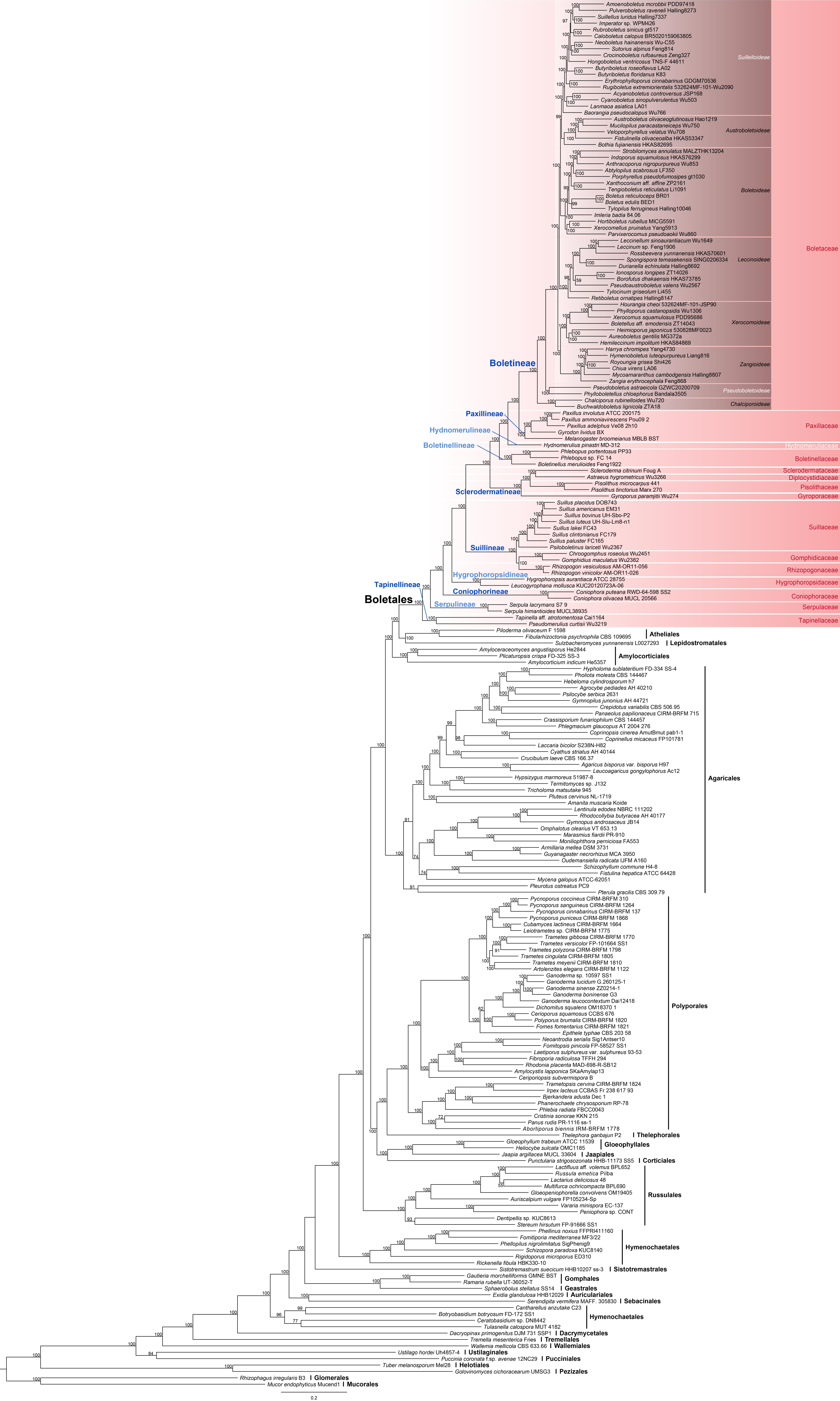

### Fig. S2. Phylogenomic_tree_0.90.jpg

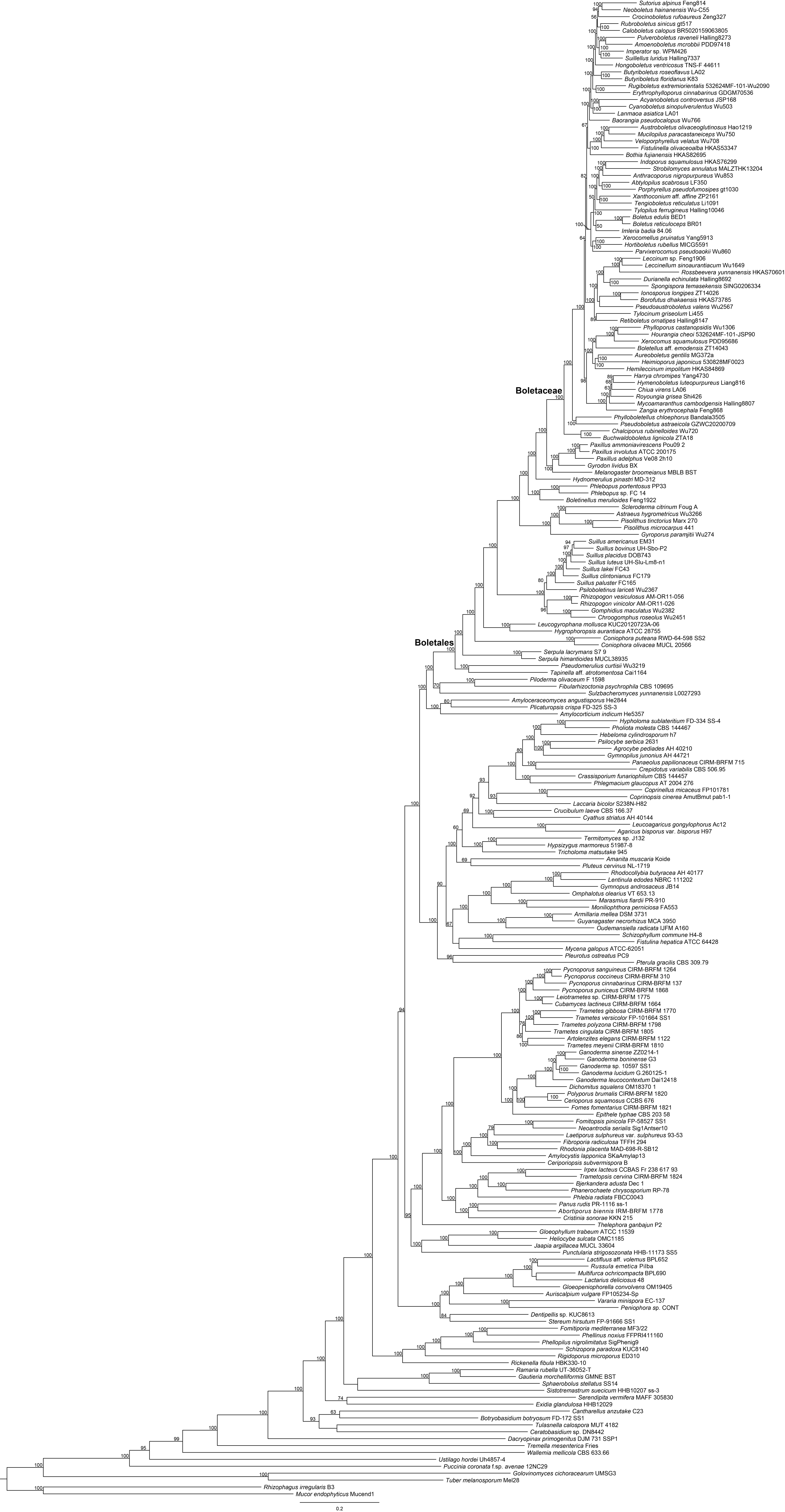

### Fig. S3. Phylogenetic_tree_four_genes.jpg

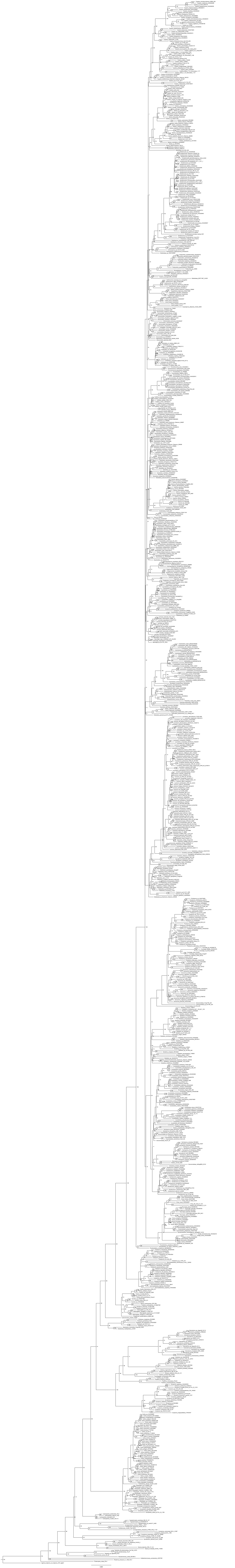

### Fig. S4. The calibrated phylogenomic tree_0.90.jpg

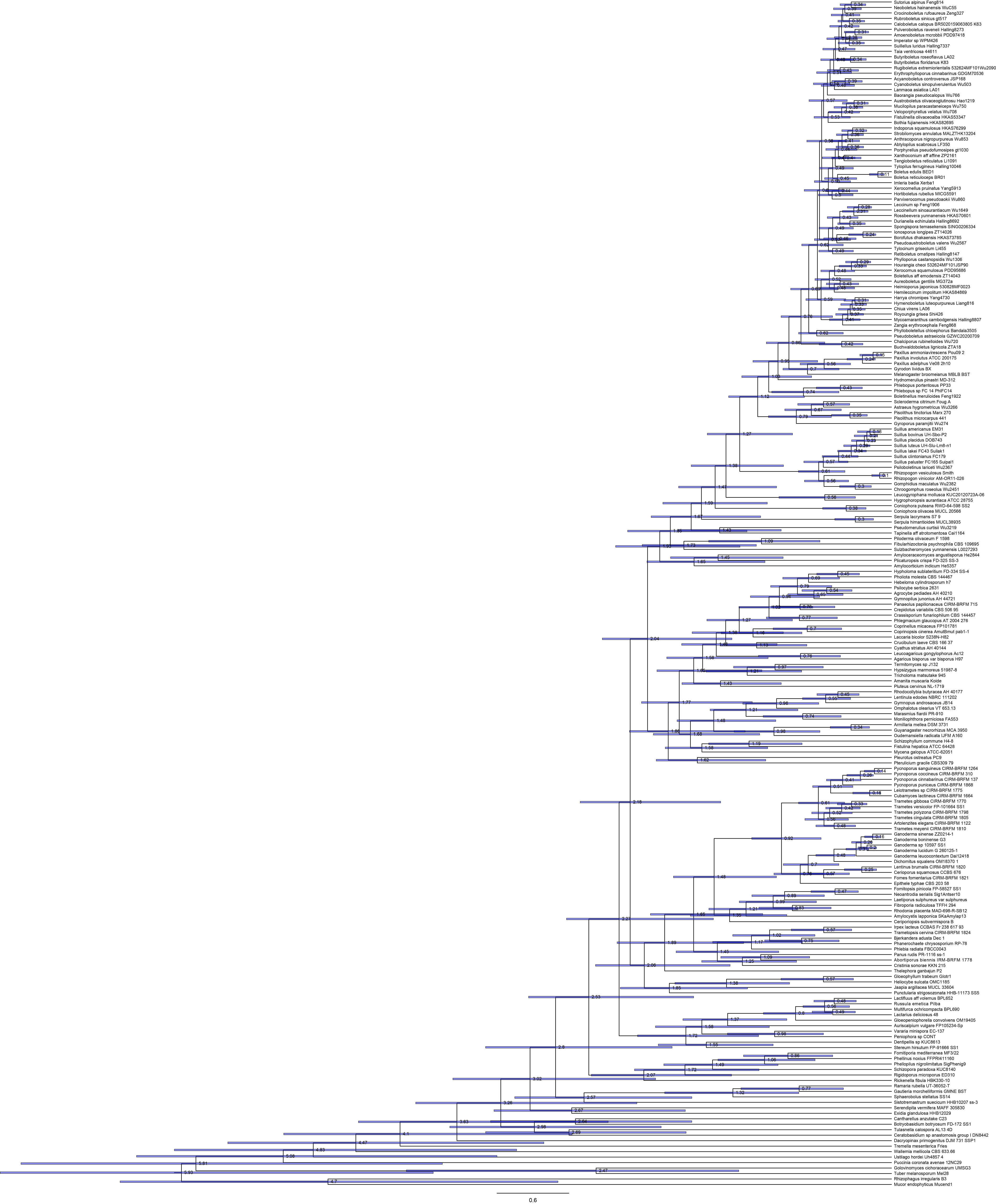

### Fig. S5. The calibrated phylogenetic tree_foru.jpg

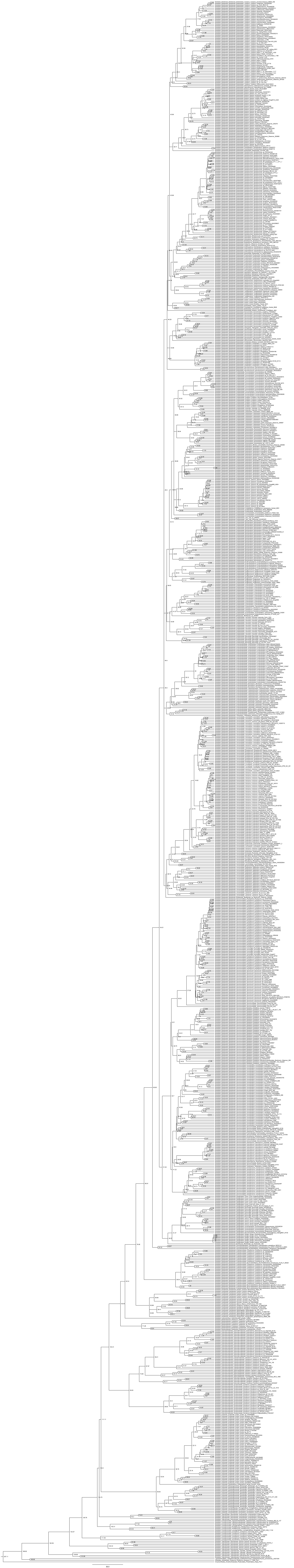

### Fig. S6. plotRateThroughTime_suborders.jpg

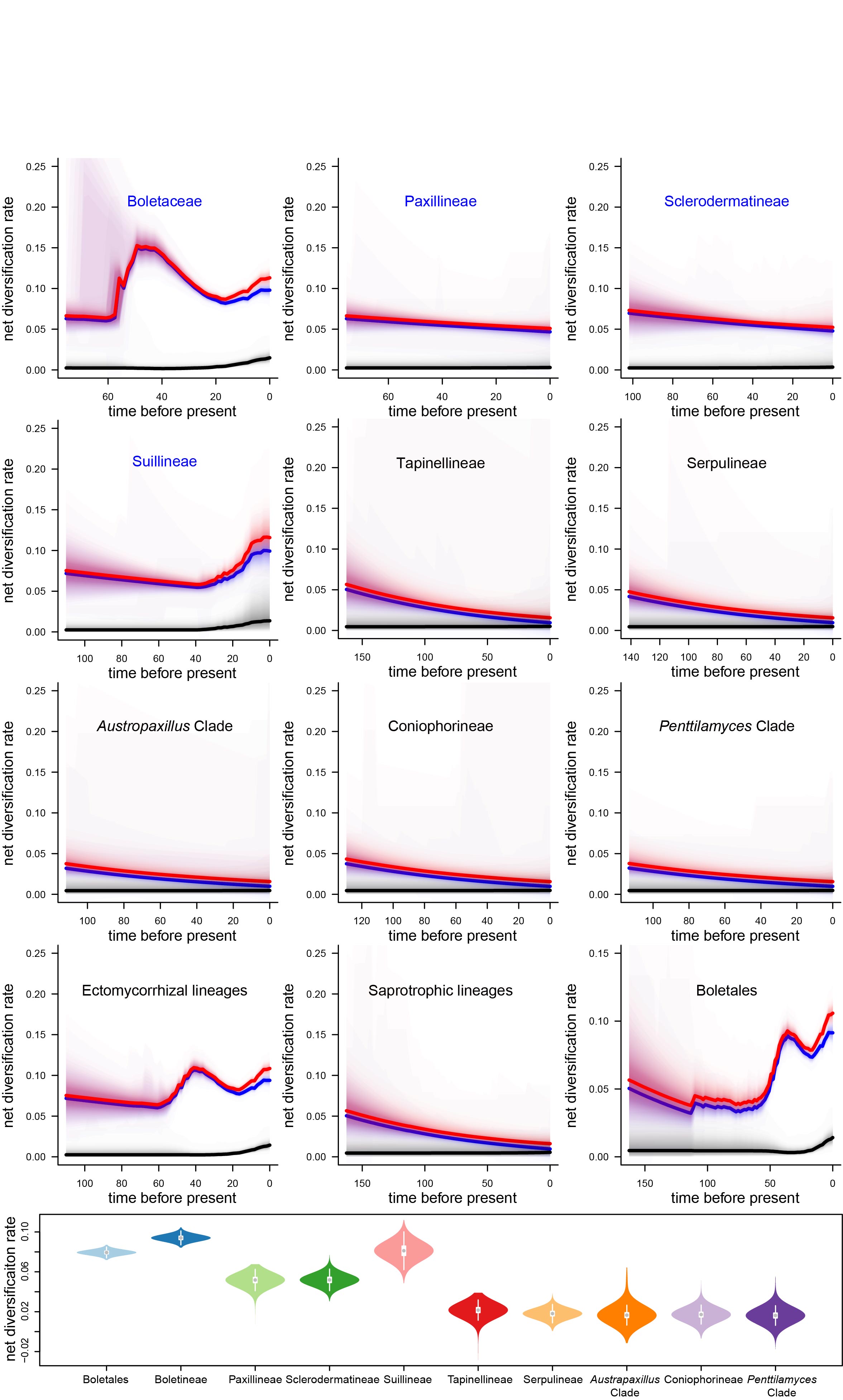

### Fig. S7. plotRateThroughTime_subfamilies.jpg

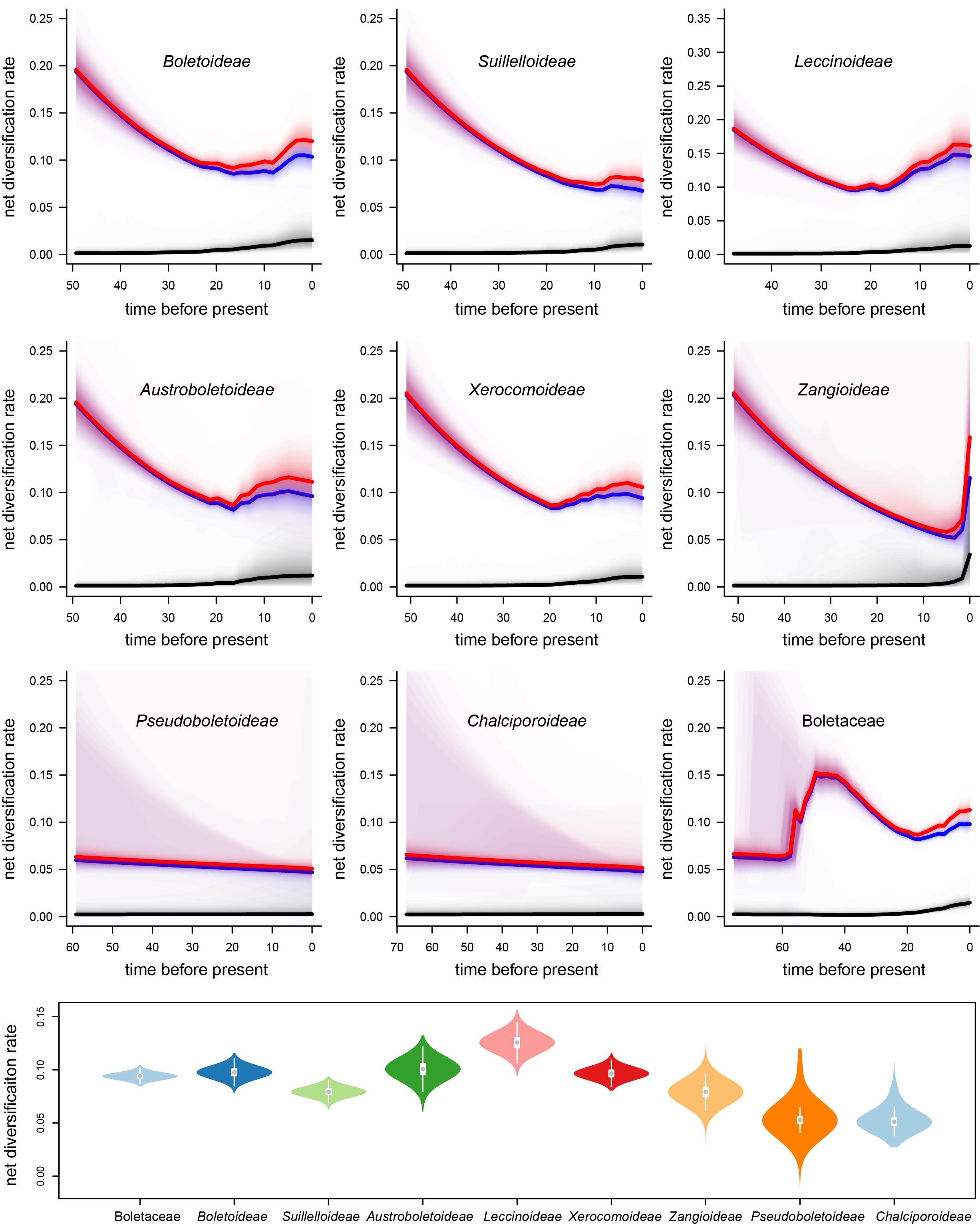

### Fig. S8. Estimated branch-specific net diversification rates.jpg

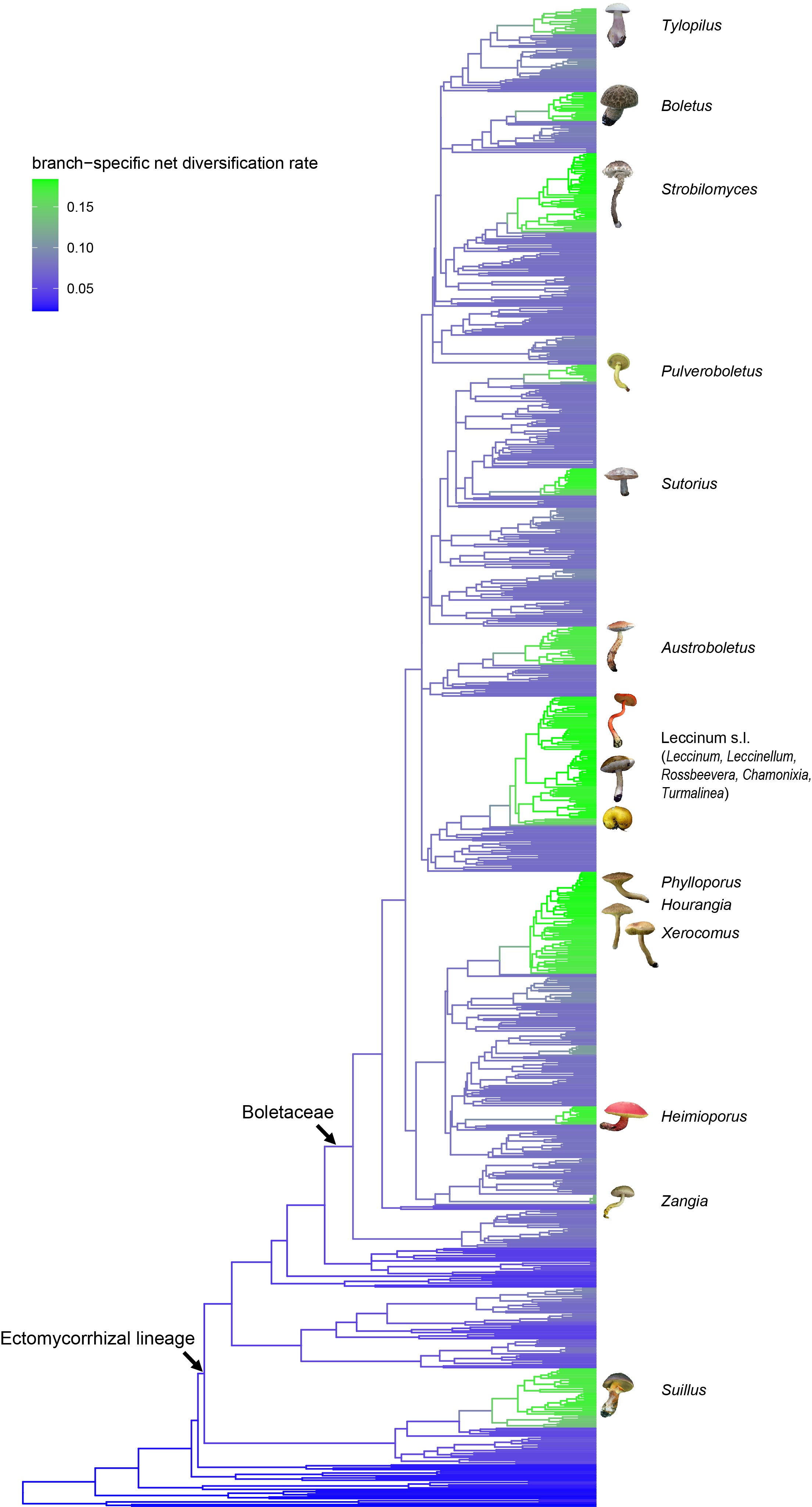

### Fig. S9. geosse_analysis_speciation+extinction+netiv.jpg

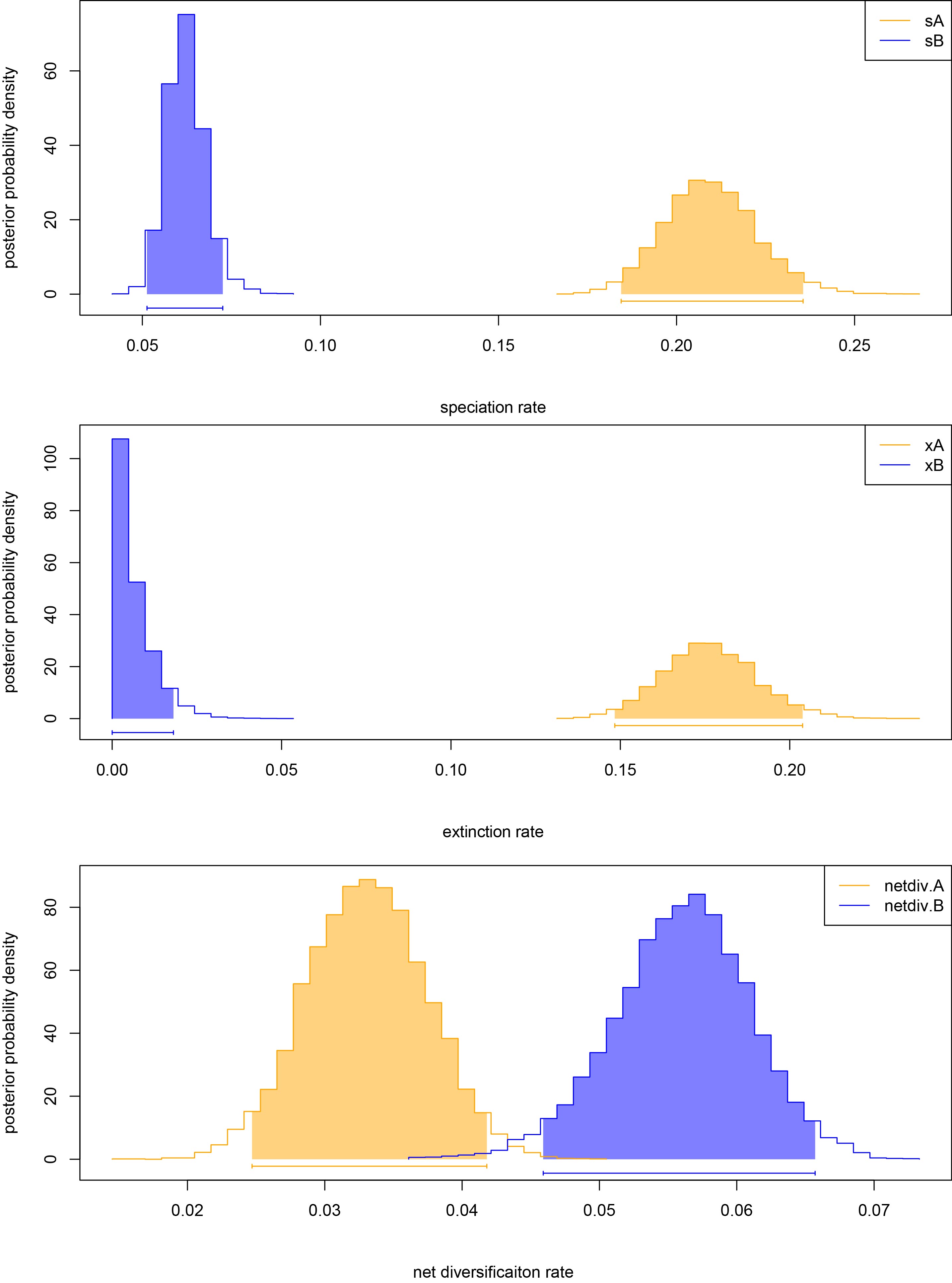

### Fig. S10. Boletales_membrane_MRCA_re.jpg

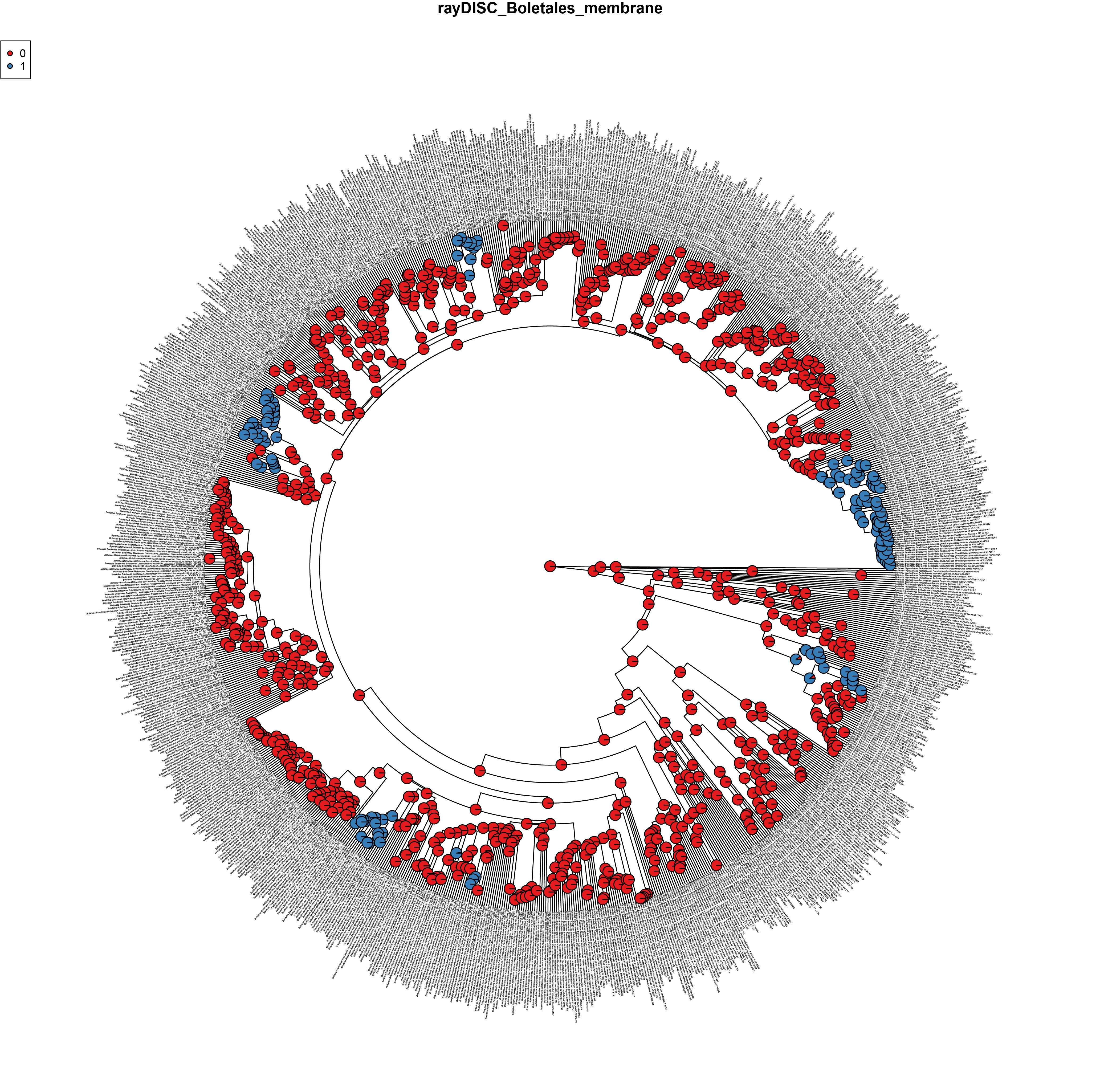

### Fig. S11. Boletales_gasteroid_MRCA.jpg

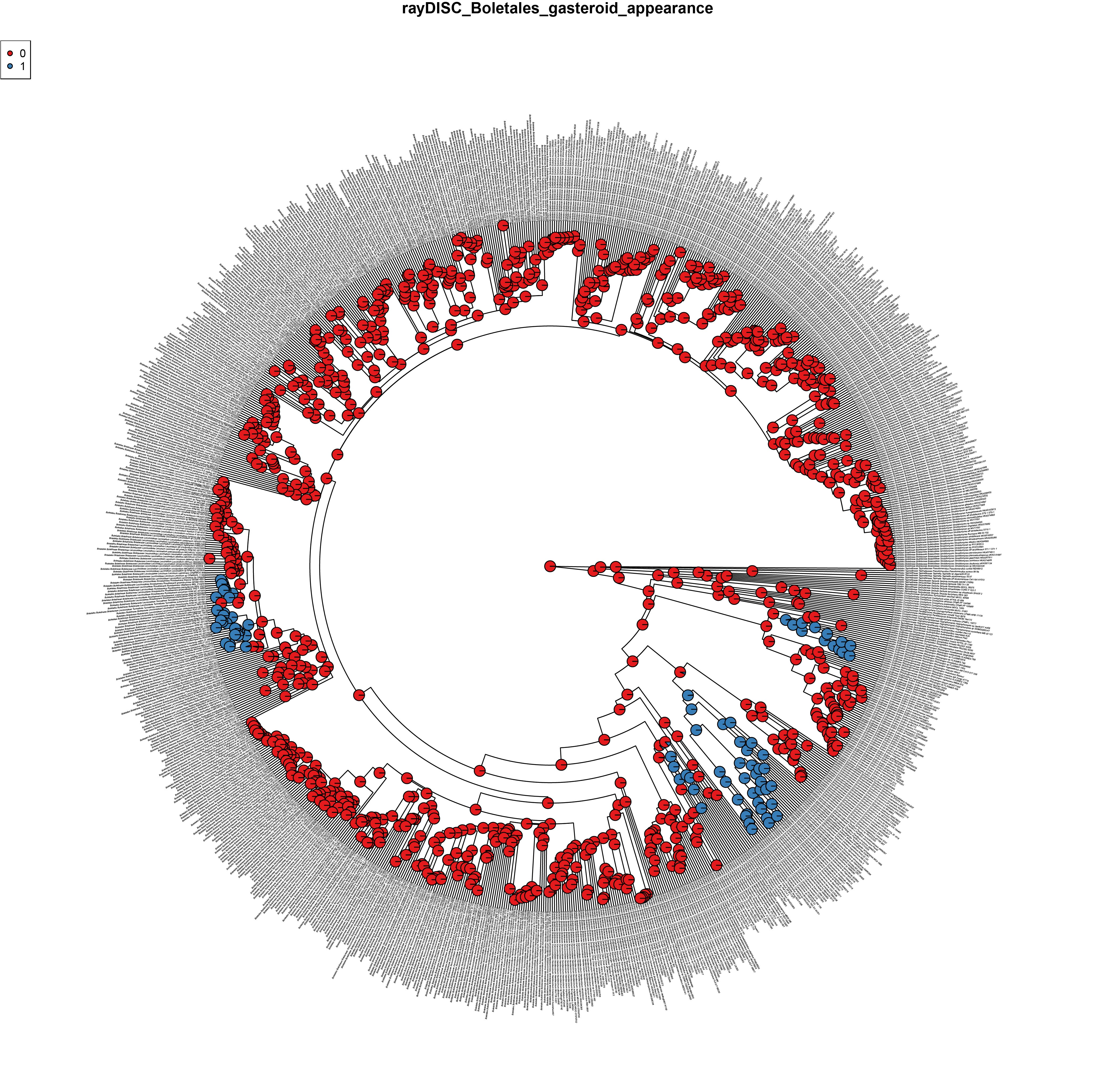

### Fig. S12. Boletales_ornamentation_MRCA.jpg

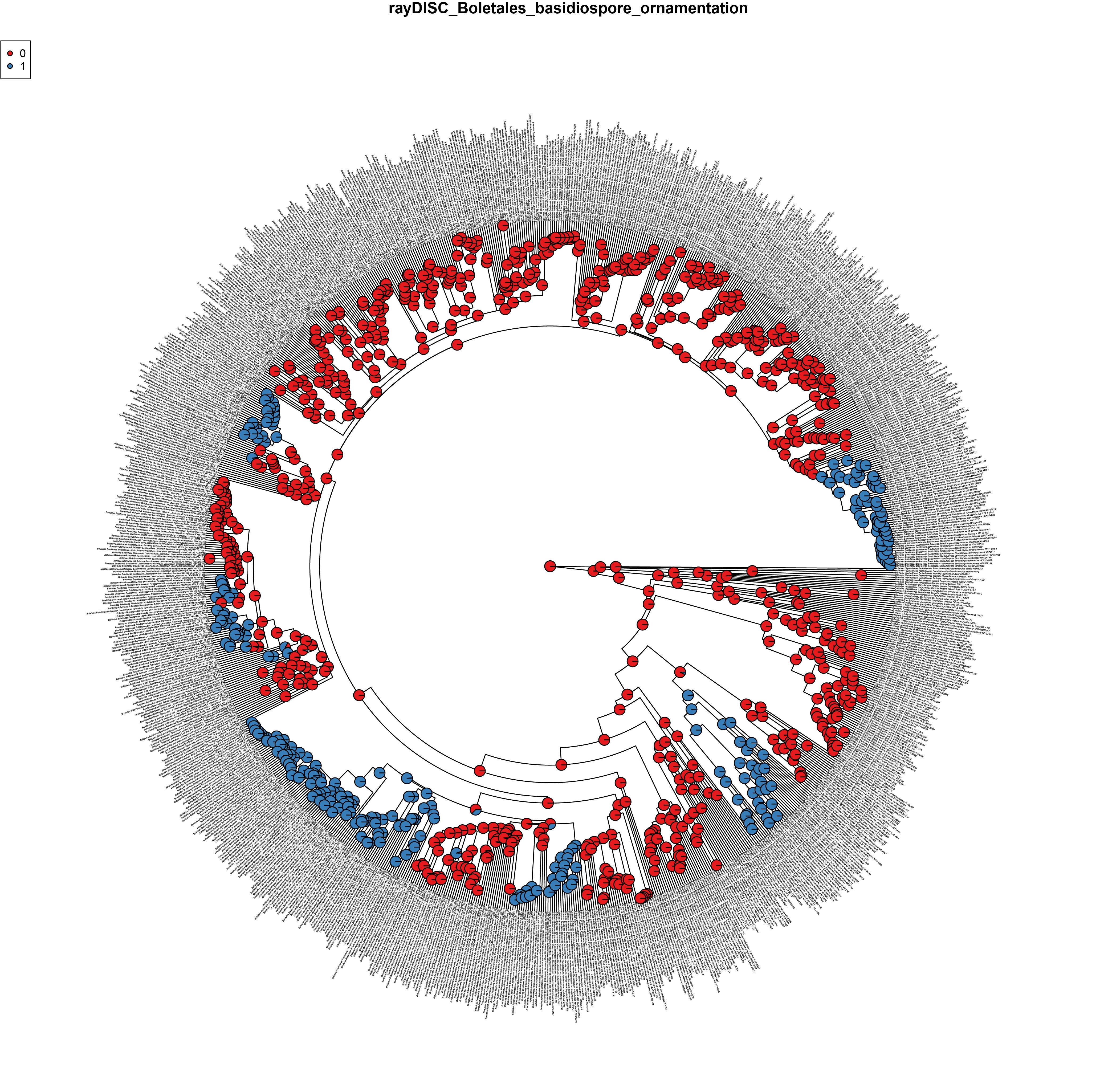
